# Rethinking human *AMY1* copy number evolution in light of demographic history

**DOI:** 10.64898/2026.02.18.706577

**Authors:** Andrea Soler i Núñez, Cécile Jolly, Camille Humbert, Afifa Chowdhury, Susanne T. Green, Pakou Harena, Lebarama Bakrobena, Forka Leypey Mathew Fomine, Peter Ebbesen, Zelalem GebreMariam Tolesa, Wendawek Abebe Mengesha, Minique de Castro, Vinet Coetzee, Himla Soodyall, Leon Mundeke, Igor Matonda, Joseph Koni Muluwa, Koen Bostoen, Sara Pacchiarotti, Johanna von Seth, Torsten Günther, Concetta Burgarella, Carina M. Schlebusch

**Affiliations:** Human Evolution, Department of Organismal Biology, Uppsala University, Uppsala, Sweden; School of Geography, Archaeology and Environmental Studies, University of the Witwatersrand; Department of History and Archaeology, University of Lomé, Togo; Department of History and African Civilizations, Faculty of Arts, University of Buea, Cameroon; Department of Health Science and Technology, University of Aalborg, Aalborg, Denmark; Department of Microbial Sciences and Genetics, College of Natural and Computational Sciences, Addis Ababa University; Department of Biochemistry, Genetics and Microbiology, University of Pretoria, Pretoria, South Africa; Division of Human Genetics, School of Pathology, Faculty of Health Sciences, University of the Witwatersrand, Johannesburg, South Africa; Academy of Science of South Africa, Pretoria, South Africa; University of Kinshasa, Kinshasa, Democratic Republic of Congo; Department of History, Ghent University, Belgium; Institut Supérieur Pédagogique de Kikwit, Kikwit, Democratic Republic of the Congo; UGent Centre for Bantu Studies, Department of Languages and Cultures, Ghent University, Belgium; SciLifeLab, Uppsala, Sweden; AGAP Institut, University of Montpellier, CIRAD, INRAE, Institut Agro, Montpellier, France; Palaeo-Research Institute, University of Johannesburg, Johannesburg, South Africa; Center for the Human Past, Department of Organismal Biology, Uppsala, Sweden

## Abstract

Dietary change is often invoked as a major selective force in recent human evolution, with increased copy number of the salivary amylase gene (*AMY1*) widely cited as an adaptation to starch-rich agricultural diets. However, most evidence for this model comes from limited geographical sampling and analyses that do not fully account for shared ancestry. Here we combine newly generated droplet digital PCR estimates from 390 individuals representing 30 Sub-Saharan African populations with published copy number data from up to 1,307 individuals worldwide and re-evaluate *AMY1* evolution using ancestry-aware and phylogenetically informed models. Across Africa, *AMY1* copy number shows no consistent association with agriculture once population structure is accounted for. At a global scale, differences between agriculturalists and non-agriculturalists are substantially smaller than previously reported and are largely explained by shared ancestry rather than diet. Phylogenetic analyses further reveal baseline differences in *AMY1* copy number between Sub-Saharan and non-Sub-Saharan populations, pointing to deep demographic processes shaping present-day variation. These results challenge the long-standing “agriculture hypothesis” and identify demographic history, rather than subsistence strategy, as the primary driver of AMY1 CN evolution worldwide.

## Introduction

Dietary change has long been considered a major selective force in human evolution, shaping both biological and cultural variation across hominin history. Key transitions, including the adoption of meat eating or scavenging, the advent of fire cooking, and the domestication of plants and animals, have left biological signatures that have traditionally been studied through archaeological artefacts, fossil morphology, and genetic variation in present-day and ancient hominins.

The adoption of agriculture and animal husbandry over the past ∼12,000 years represents the most recent major shift in human diets. This transition led to a significant reduction in food diversity, as agriculturalist societies relied on fewer domesticated plant and animal species (Armelagos, 2014). Dietary starch consumption increased, and it accounts for up to 70% of the caloric intake of modern agricultural diets (Copeland et al., 2009). In humans, the digestion of this complex polysaccharide is initiated in the oral cavity by salivary amylase (AMY1) and continues in the small intestine through the action of pancreatic amylases (AMY2A, AMY2B) and other enzymes (e.g., MGAM, SI) until its complete breakdown into glucose molecules (Pedersen et al., 2002).

Diversity at the amylase locus is not reflected at the nucleotide level but rather in its extensive copy number (CN) variation, with diploid *AMY1* CN ranging from two to twenty among present-day humans (Bolognini et al., 2024). Several studies, largely based on Eurasian populations, have reported positive correlations between *AMY1* CN and amylase activity in the saliva or serum (Perry et al., 2007; Mandel et al., 2010; Yang et al., 2014; Tan et al., 2015; Carpenter et al., 2017; Atkinson et al., 2018; Nayema et al., 2023). Adding to this findings, two population-level analyses further reported higher *AMY1* CN in groups that have traditionally consumed high-starch or agricultural diets compared to those practicing other subsistence strategies (Perry et al., 2007; Bolognini et al., 2024).

Nevertheless, the evolutionary history of the *AMY1* locus predates the emergence of agriculture and extends beyond modern humans. Comparative analyses indicate that present-day human *AMY1* haplotypes share a most recent common ancestor after the divergence from Neandertals and Denisovans, with a three-copy haplotype seeding much of the variation observed today (Inchley et al., 2016; Bolognini et al., 2024; Yilmaz et al., 2024). Analyses of archaic genomes suggest that the ancestral state of the locus was likely a single-copy haplotype, although higher *AMY1* CN has been detected in at least one Denisovan and several Neandertal individuals, indicating that CN polymorphism was already present in archaic hominin lineages (Prüfer et al., 2014; Perry et al., 2015; Inchley et al., 2016; Bolognini et al., 2024; Yilmaz et al., 2024).

Ancient DNA studies focusing on West Eurasia have reported an increase in total CN of amylase genes over the past 12,000 years, coinciding with the spread of agriculture in Europe, and have suggested moderate selective pressures favouring high CN during this period (Bolognini et al., 2024; Yilmaz et al., 2024). However, high *AMY1* CN have also been observed in 45,000 and 34,000 year-old individuals, reinforcing the view that substantial variation in this locus existed well before the onset of agriculture in the region (Yilmaz et al., 2024).

Despite recent advances in understanding the worldwide diversity and haplotype structure of the amylase locus (Bolognini et al., 2024; Yilmaz et al., 2024), associations between *AMY1* CN and dietary practices are based on limited datasets that do not represent the genetic and cultural diversity of populations across continents (Perry et al., 2007; Bolognini et al., 2024). Here, we address this gap by generating new droplet digital PCR (ddPCR) estimates of *AMY1* CN for 390 individuals from 30 Sub-Saharan African populations spanning diverse ancestries and subsistence strategies (Figure 1A). We combine these data with published *AMY1* CN estimates from global population panels, comprising up to 1,307 individuals in total (Bolognini et al., 2024) (Figure 1B), and re-evaluate the evolution of this gene using ancestry-aware and phylogenetically informed models. This framework allows us to disentangle dietary and demographic contributions to global patterns of *AMY1* CN variation.

**Figure 1.**
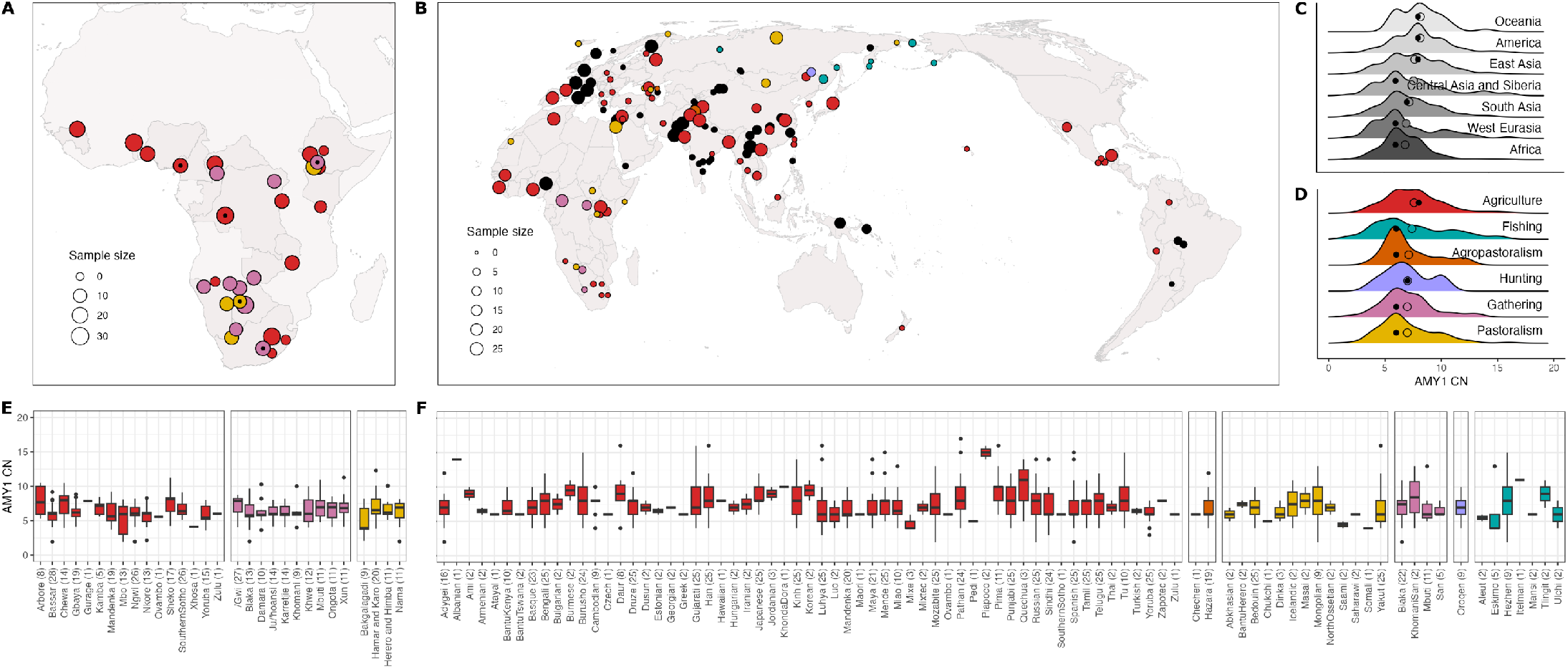
Geographical location of the populations in the African **ddPCR data set** (n=390, N=30) (**A**) and the complete **read-depth data set** (n=1307, N=136) (**B**), and their *AMY1* copy number (CN) estimates (**C**-**F**). Populations in all panels are colored by their dominant subsistence strategy (agriculture, pastoralism, agropastoralism, hunting, gathering, fishing). All colored populations are included in the modeling framework: colored circles with a black dot in the first panel (**A**) represent low-annotation groups in the **ddPCR data set** for which only the main subsistence was annotated. Black circles in the second panel (**B**) represent groups in the **read-depth data set** for which subsistence is not documented and are therefore not included in the modeling framework but only the phylogenetic tests. Density plots in panels **C** and **D** show *AMY1* CN distributions in the **read-depth data set** by region (**C**) and subsistence strategy (**D**). Full and empty circles represent the median and mean *AMY1* CN for each plotted group, respectively. Groups are sorted by their mean.

## Results

### No support for an effect of agriculture on *AMY1* CN variation in Africa

To assess whether variation in *AMY1* CN is associated with subsistence strategy in Africa, we analyzed both a newly generated ddPCR dataset and a read-depth dataset using ancestry-aware regression models that account for shared genetic background among individuals and populations. Both Bayesian and frequentist modeling approaches were used, since they allow to correct for shared ancestry in complementary ways (see Table 1, Methods and Supplementary Methods). Across all analyses, *AMY1* CN showed no consistent association with agriculture among African populations.

**Table 1.** Model specifications.

| Model | Random effects |
| --- | --- |
| All models | Individual-level **phylogenetic VCV** (only in *brms* framework). |
| All models | **Population identity** (discrete), e.g. 'Kemba', 'Mbuti', 'Gbayá', 'Yakut'. |
| Model | Fixed effects |
| All models | Genetic **principal components** (only in *glmmTMB* framework). |
| Agr.vs.NonAgr | A group's **binary subsistence category** (discrete): 'Agriculture' or 'Non-Agriculture'. |
| **All.vs.All** | A group's **multi-level subsistence category** (discrete): 'Agriculture', 'Pastoralism', 'Hunting', 'Gathering', 'Fishing' or, less often, 'Agropastoralism'. |
| **Agriculture** | A group's relative **dependence on agriculture** (continuous-like, ILR-transformed). |
| **Pastoralism** | A group's relative **dependence on pastoralism** (continuous-like, ILR-transformed). |
| **Hunting** | A group's relative **dependence on hunting** (continuous-like, ILR-transformed). |
| **Gathering** | A group's relative **dependence on gathering** (continuous-like, ILR-transformed). |
| **Fishing** | A group's relative **dependence on fishing** (continuous-like, ILR-transformed). |

In the complete African ddPCR dataset (n=390 individuals from N=30 Sub-Saharan populations), *AMY1* CN was not associated with agriculture whether subsistence was modeled as a binary category (agriculturalist vs. non-agriculturalist), a multi-level categorical variable (agriculturalist, pastoralist, hunter, gatherer, fisher), or a continuous measure of dependence on agriculture. This result was consistent when restricting analyses to the subset of highly annotated groups (n=317, N=25) and agriculturalists (n=168, N=14). In all cases, effect estimates for agriculture were centered around zero, and model comparisons consistently favored a null model that accounted only for genetic ancestry (Figures 2A-2C; Tables S3.1-S3.3). Analyses of African populations in the read-depth dataset (n=163, N=20) yielded similar results. When subsistence variables were added to models accounting for population structure, agriculture did not explain additional variation in *AMY1* CN (Tables S3.4 and S3.5). These findings indicate that, across Africa, subsistence strategy does not constitute a robust predictor of *AMY1* CN once population structure is taken into account.

**Figure 2.**
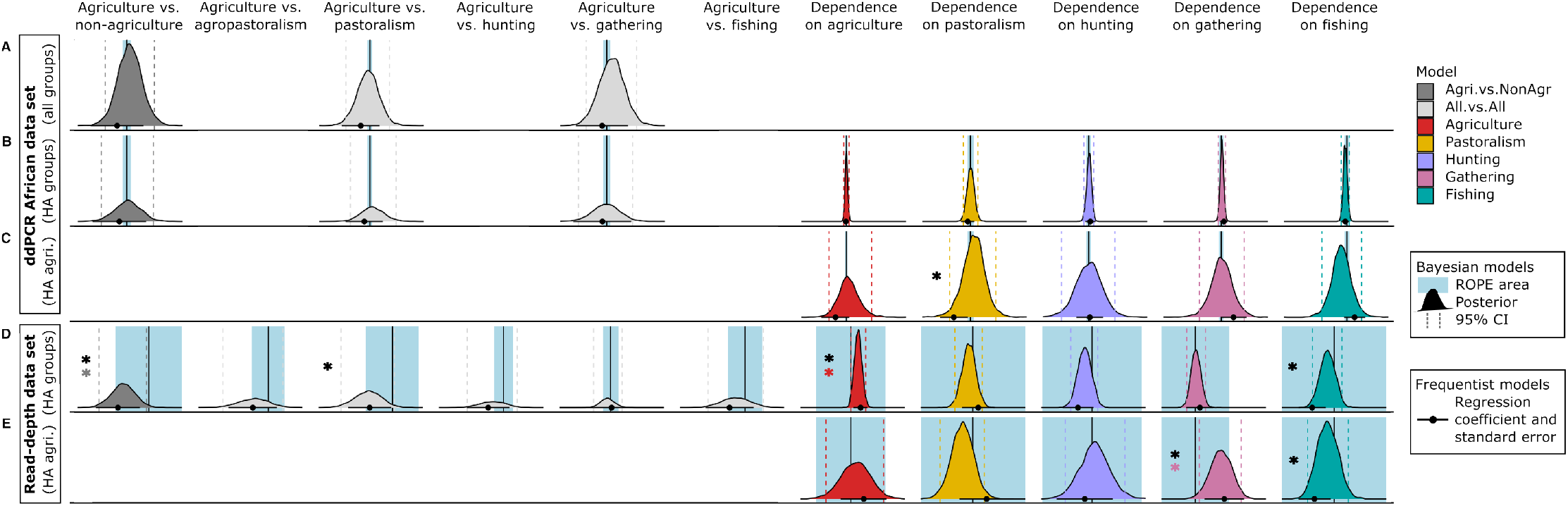
Results from the Bayesian and frequentist models on the **ddPCR** (**A**-**C**) and **read-depth** (**D**-**E**) data sets. Plot grids are separated row-wise to describe different data sets, while columns refer to the effect of variables evaluated in a given model and data set. The effect of all variables is plotted as follows: (i) posterior distributions (Bayesian models) and 95% CI around the posterior mean are represented as colored density plots and dashed lines, while (ii) regression coefficients and their corresponding standard error (frequentist models) are drawn as black dots and lines at the bottom of each grid. Blue shaded areas at the back of each grid represent standard ROPE ranges (−0.1, 0.1) and black vertical lines mark the zero on the x axis. Colored and black asterisks on each grid indicate that the effect of the variable in a given data set and model is significant, by the Bayesian and/or frequentists models, respectively. Note that the scale of the x axis is only shared among grids of the same data set (**A**-**C** and **D**-**E**).

The only statistically supported associations with subsistence practices in Africa were detected under the frequentist modeling framework. In this context, dependence on gathering showed a positive association with *AMY1* CN in the read-depth dataset (n=163, N=20) and significantly improved model fit relative to the corresponding null model. This association persisted when analyses were restricted to agriculturalists alone (n=113, N=11). In contrast, in the ddPCR dataset restricted to agriculturalist populations (n=168, N=14), dependence on gathering showed a similar effect but did not reach statistical significance, whereas dependence on pastoralism was negatively associated with *AMY1* CN and improved model fit (Tables S3.2-S3.5).

### Genetic ancestry is the primary predictor of *AMY1* CN variation in Africa

In contrast to subsistence-related variables, genetic ancestry and population structure consistently emerged as significant predictors of *AMY1* CN variation among African populations. This pattern was observed across all models of the ddPCR dataset (Tables S3.1-S3.3).

In ancestry-aware regression models controlling for population structure, genome-wide principal components explaining genetic variation accounted for a substantial fraction of the observed differences in *AMY1* CN. In particular, the principal component capturing Eurasian-related ancestry was positively associated with *AMY1* CN across all models and data partitions, indicating that African individuals with higher proportions of Eurasian ancestry (most prominently, Ethiopian populations) tend to exhibit higher *AMY1* CN (Figure 3B).

**Figure 3.**
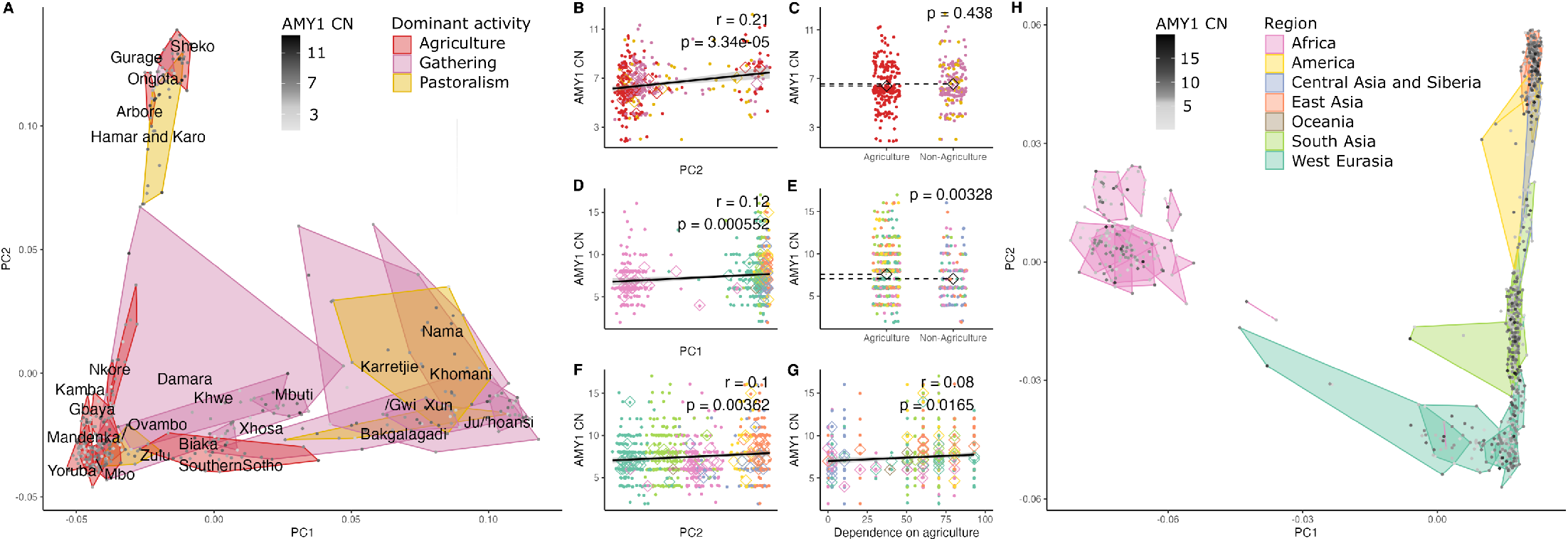
*AMY1* copy number (CN) and principal components analysis (PCA) of individuals from the complete African **ddPCR data set** (n=390, N=30) (**A**-**C**) and the highly annotated groups in the **read-depth data set** (n=810, N=68) (**D**-**H**). Both PCA plots are based on SNP markers on chromosome one. Each individual is represented by a dot colored by its *AMY1* CN, and each population is enclosed by a polygon colored by its dominant subsistence strategy (**A**) or geographical region (**H**). The remaining plots (**B**-**G**) show the relationship between *AMY1* CN and principal components or subsistence variables in the two data sets. Correlation (r) and p-values (p) in plots **B, D, F** and **G** refer to a Pearson’s correlation test, and the trend line represents an isolated linear model between the two plotted variables. Additionally, population averages are specified as colored rhombi. In plots **C** and **E**, p-values (p) correspond to a two-sided Wilcoxon test between agriculturalists and non-agriculturalists in the two data sets, and black rhombi represent their average *AMY1* CN.

These findings were supported by Bayesian models explicitly incorporating population identity and individual-level phylogenetic structure. Across all model formulations and data subsets, both population-level effects and phylogenetic relatedness explained significant variation in *AMY1* CN, with credible intervals excluding zero (Tables S3.1-S3.3). Notably, the phylogenetic component remained well supported despite the use of a conservative prior, underscoring the robustness of ancestry effects at the individual level.

Together, these results indicate that population structure and shared genetic ancestry, rather than subsistence strategy, constitute the primary determinants of *AMY1* CN variation within Africa.

### Ancestry and diet jointly shape *AMY1* CN variation at a global scale

We next extended our analyses to a global scale using the read-depth dataset, which includes 810 individuals from 86 populations with detailed subsistence annotations. In this worldwide context, both modeling frameworks detected a modest but statistically supported difference in *AMY1* CN between agriculturalist and non-agriculturalist populations, as well as a positive association between *AMY1* CN and dependence on agriculture (Figures 2D, 3E and 3G; Table S3.6).

The magnitude of this difference was small: agriculturalist populations exhibited, on average, 0.53 additional diploid copies of *AMY1* compared to non-agriculturalists (mean *AMY1* CN = 7.58 vs. 7.05). Among individual subsistence contrasts, only the comparison between agriculturalists and pastoralists was significant across modeling approaches, while hunters, gatherers, fishers, and agropastoralists did not differ from agriculturalists in *AMY1* CN. Dependence on fishing was negatively associated with *AMY1* CN under the frequentist framework, and models including fishing, agriculture, or the agriculturalist vs. non- agriculturalist contrast modestly improved model fit relative to the null model.

Despite these statistically supported associations, Bayesian effect-size assessments indicated that the contribution of agriculture was small. The posterior distributions for both dependence on agriculture and the agriculturalist vs. non-agriculturalist contrast were largely compatible with negligible effects (Figure 2D; Table S3.6), indicating that diet explains only a limited fraction of global *AMY1* CN variation once ancestry is accounted for.

Across all global models, genetic ancestry remained a strong and consistent predictor of *AMY1* CN. Both the individual-level phylogenetic structure in the Bayesian framework and principal components summarizing genetic ancestry in the frequentist framework showed non-zero effects across nearly all models (Table S3.6). In particular, *AMY1* CN was higher in individuals with reduced African ancestry and increased East Asian or American ancestry components (Figures 3D and 3F).

When analyses were restricted to agriculturalist populations alone (n=640, N=59), dependence on agriculture no longer predicted *AMY1* CN. Instead, dependence on gathering showed a positive association with *AMY1* CN across modeling frameworks, while dependence on fishing was negatively associated under the frequentist approach (Figure 2E; Table S3.7). As in the full global analysis, posterior distribution assessments indicated that the effect of gathering was small, with most of the posterior distribution falling within the region of practical equivalence (Figure 2E).

Taken together, these results show that while subsistence-related signals can be detected at a global scale, their effect sizes are modest, context-dependent, and are consistently outweighed by the contribution of shared genetic ancestry to *AMY1* CN variation.

### Phylogenetic analyses reveal distinct baseline *AMY1* CN distributions between Sub-Saharan and non-Sub-Saharan populations

To evaluate whether *AMY1* CN variation reflects deeper demographic structure beyond subsistence practices, we tested for phylogenetic signal across the human population tree using three complementary approaches: global statistics of phylogenetic signal, phylogenetic correlograms, and local indicators of phylogenetic association (LIPA).

Across both the ddPCR and read-depth datasets, measures of global phylogenetic signal detected a weak but significant departure from the null model, indicating that closely related individuals tend to exhibit more similar *AMY1* CN values than expected by chance (Tables S4.1 and S4.2). Phylogenetic correlogram tests added detail to the previous results by showing that the phylogenetic signal is concentrated at short distances and not present across all clades of the human phylogeny. Specifically, we found low but significant positive autocorrelation among closely related individuals, which rapidly decayed with increasing phylogenetic distance and was no longer detectable beyond 0.25 units of phylogenetic distance (Figures 4D-4H). This pattern was consistent across all non-African population subsets of the read-depth dataset, including West Eurasia and South Asia (n=712, N=61), East and Central Asia, Siberia, Oceania, and the Americas (n=407, N=53), and all groups from the previous regions (n=1119, N=115). In contrast, Sub-Saharan African populations deviated from this pattern. No significant phylogenetic autocorrelation was detected at any phylogenetic distance within Africa in either the ddPCR (Figure S4) or read-depth datasets.

**Figure 4.**
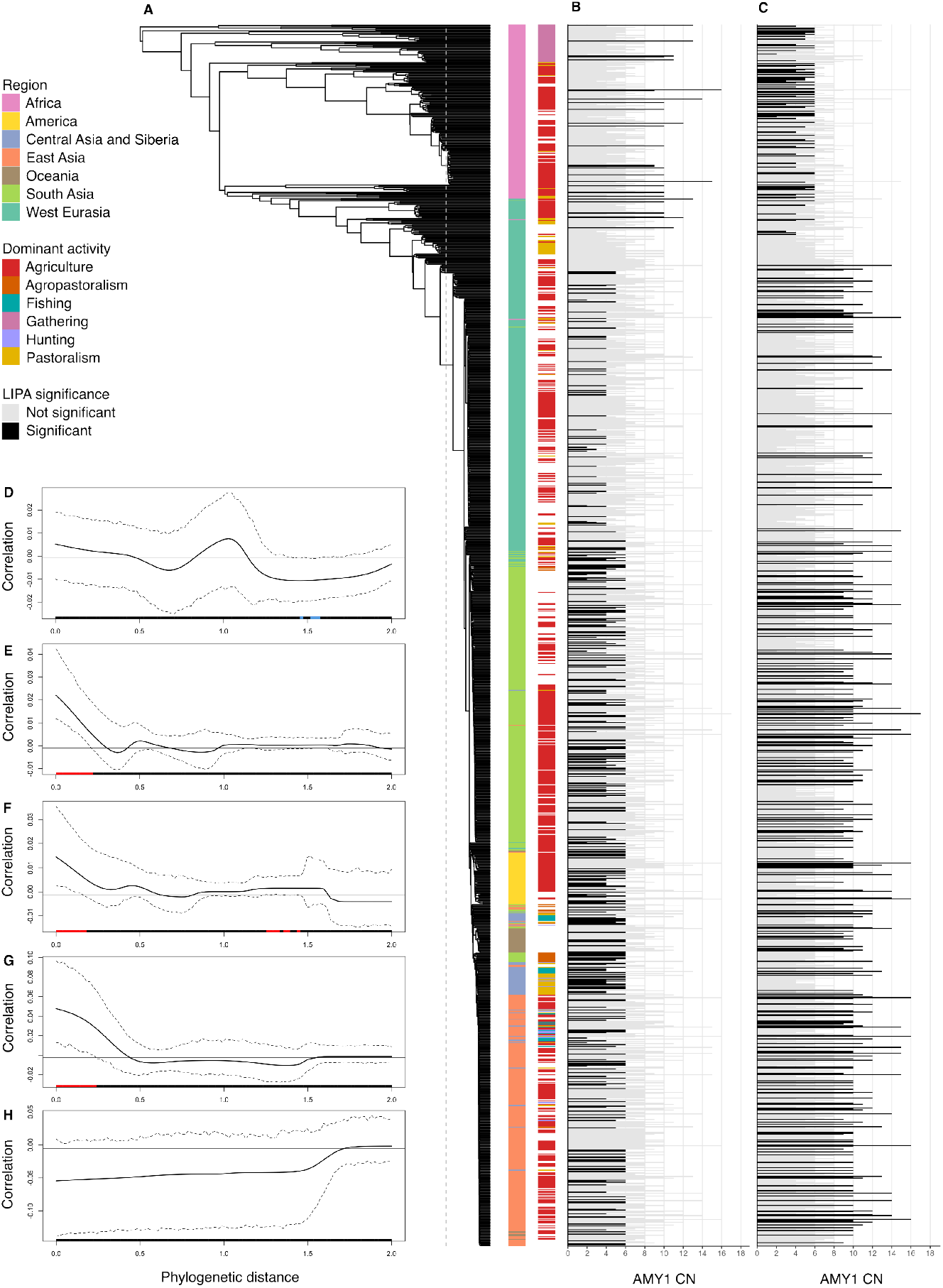
Phylogenetic tree of the **read-depth data set** (n=1307, N=135) (**A**), with results from the local indicators of phylogenetic association (LIPA) tests (**B, C**) and phylogenetic correlograms (**D, E, F, G, H**). Each terminal branch in the phylogenetic tree (**A**) corresponds to an individual in the **read-depth data set** and the two associated colored plots correspond to the geographical region (left) and dominant activity (right) of the population to which each individual belongs. A gray dashed line marks one eight of phylogenetic distance from the tips to the root. Individual *AMY1* CN and results of the LIPA test for the ‘less’ and ‘greater’ hypotheses are shown in the second (**B**) and third (**C**) panels, respectively. Each bar is colored by the significance of its LIPA test (alpha=0.05). The remaining panels show results for the phylogenetic correlogram tests using the complete **read-depth data set** (n=1307, N=136) (**D**), the non- Sub- Saharan populations (n=1119, N=114) (**E**), the West Eurasian and South Asian populations (n=712, N=61) (**F**), the East and Central Asian, Siberian, Oceanian and America populations (n=407, N=53) (**G**), and the Sub-Saharan populations (n=188, N=21) (**H**). Red and blue colors indicate significant positive and negative correlations, respectively, at the indicated phylogenetic distance.

LIPA tests further revealed contrasting baseline distributions of *AMY1* CN between Sub-Saharan and non-Sub-Saharan clades. Under the ‘greater’ hypothesis, which detects clusters of similar trait values, LIPA identified significant positive autocorrelation in the non-African and Ethiopian clades, which were composed of individuals with high *AMY1* CN (Figure 4C). Conversely, under the ‘less’ hypothesis, which identifies outlier trait values, significant LIPA measures were observed in non-African and Ethiopian individuals with low *AMY1* CN, as well as a few Sub-Saharan individuals with unusually high *AMY1* CN (Figure 4B). This reciprocal pattern was consistently observed across both the ddPCR (Figure S4) and the read-depth datasets.

Together, these results indicate that *AMY1* CN variation is structured around different baseline distributions in Sub-Saharan (lower *AMY1* CN) and non-Sub-Saharan (higher *AMY1* CN) populations, consistent with deep demographic processes rather than recent, localized adaptation.

### Regional patterns of *AMY1* CN variation outside Sub-Saharan Africa

Using the size of the read-depth data set, we next examined whether regional patterns outside Sub-Saharan Africa mirror the global associations between *AMY1* CN, subsistence strategy, and ancestry. We separately modeled *AMY1* CN in populations from (i) West Eurasia and South Asia, and (ii) East and Central Asia, Siberia, Oceania, and the Americas.

Among West Eurasian and South Asian populations, *AMY1* CN showed no robust association with subsistence strategy. Across models, subsistence-related predictors failed to explain variation in *AMY1* CN, with only a marginal positive association between *AMY1* CN and dependence on gathering detected among agriculturalist populations (n=358, N=26; Tables S3.8 and S3.9).

In contrast, populations from East and Central Asia, Siberia, Oceania, and the Americas broadly recapitulated the global pattern. In analyses including all populations from these regions (n=236, N=33), *AMY1* CN was higher in agriculturalists than non-agriculturalists (mean *AMY1* CN = 8.28 vs. 7.21) and increased with dependence on agriculture (Tables S3.10 and S3.11). However, Bayesian effect-size assessments indicated that the contribution of agriculture was small: reliance on agriculture showed a negligible effect, with its entire posterior distribution falling within the region of practical equivalence, whereas the categorical distinction between agriculturalists and non-agriculturalists showed partial support. When analyses were restricted to agriculturalist populations alone (n=169, N=22), neither modeling framework detected an effect of subsistence on *AMY1* CN.

To further explore demographic influences, we examined the relationship between *AMY1* CN diversity and geographic distance from East Africa in all non-Sub-Saharan populations with at least five sampled individuals. Two measures of CN diversity (Shannon entropy and the number of unique CN values) were negatively correlated with distance from East Africa, consistent with a reduction in genetic diversity away from Africa (Figure S5).

Finally, focusing on West Eurasia, we re-examined the reported increase in *AMY1* CN over the past 12,000 years using ancient European genomes. Among 533 individuals, *AMY1* CN was negatively correlated with sample age, describing how more ancient samples have lower *AMY1* CN than more recent ones. Additionally, *AMY1* CN also showed significant associations with ancestry components related to Western hunter-gatherers and Anatolian farmers (Figure S3; Table S5). The significance of these associations was nevertheless not maintained when modeling *AMY1* CN by each ancestry component jointly with sample age, indicating that sample age is a better predictor of *AMY1* CN than any of the ancestry components alone.

## Discussion

### Demography, not agriculture, dominates global *AMY1* CN variation

This study provides the first ancestry-aware analysis of *AMY1* CN variation across Africa and worldwide. By integrating newly generated ddPCR data from diverse Sub-Saharan African populations with read-depth data to form the largest dataset assembled to date, we show that demographic history, rather than subsistence strategy, is the primary driver of present-day variation at the salivary amylase locus.

### Limits of a universal agriculture-based model

Higher *AMY1* CN has long been interpreted as an adaptation to starch-rich agricultural diets, based on comparisons between agriculturalist and non-agriculturalist populations (Perry et al., 2007; Bolognini et al., 2024). Our results challenge the universality of this model. Across Sub-Saharan Africa and West and South Eurasia, *AMY1* CN shows no consistent association with agriculture once population structure is taken into account. A modest association emerges only when populations are pooled globally, and even then the magnitude of the difference between agriculturalists and non-agriculturalists is substantially smaller than previously reported.

Notably, the estimated difference has declined from 1.28 diploid copies in early studies sampling seven populations (Perry et al., 2007), to ∼1.0 in analyses of 33 populations (Bolognini et al., 2024), to 0.53 diploid copies in the present study encompassing 86 populations. This trend suggests that earlier estimates were inflated by limited sampling and incomplete control for ancestry. In this context, the biological relevance of such small average differences becomes difficult to reconcile with the high intra-population variability of *AMY1* CN and the modest proportion of enzymatic activity it explains (Fernández & Wiley, 2017).

Comparative studies in domesticated species demonstrate that amylase gene expansions can be adaptive in strict carnivores transitioning to starch-rich diets, such as dogs (Axelsson et al., 2013), but often show weaker or absent effects in omnivorous species with ancestral starch consumption (Pajic et al., 2019). Humans, whose pre-agricultural diets frequently included substantial plant and starch intake (Eaton & Konner, 1985; Cordain et al., 2000; Hatley & Kappelman, 1980; Chen et al., 2024), may therefore not conform to a simple agricultural-adaptation model. In Sub-Saharan Africa, linguistic and ethnobotanical evidence further indicates long-standing consumption of starchy tubers prior to the adoption of intensive farming (Bostoen & Muluwa, 2017; Bostoen et al., 2025; Scarcelli et al., 2019).

### Beyond starch digestion

Although *AMY1* initiates starch digestion in the oral cavity, much of starch breakdown and glucose uptake occurs downstream through other enzymes, several of which show stronger signatures of selection (Lindo et al., 2018). Moreover, amylase is expressed across a wide range of human tissues and has been implicated in diverse physiological processes, including stress responses, immunity, reproduction, microbiome composition, and oral health (Poole et al., 2019; Kamitaki et al., 2025, Nater & Rohleder, 2009; Fuentes et al., 2011; Behringer et al., 2011; Fuentes-Rubio et al., 2016, Esterhuizen et al., 1995; Singh, 1995). The inconsistent association between *AMY1* CN and agriculture observed here, together with these broader functional roles, suggests that any selective pressures acting on this locus may extend well beyond dietary starch metabolism.

### Neutral processes and deep demographic structure

Across African and global analyses, genetic ancestry consistently explained a substantial fraction of *AMY1* CN variation. Individuals with reduced Sub-Saharan African ancestry and increased Eurasian, East Asian, or American ancestry tend to exhibit higher *AMY1* CN, a pattern reinforced by phylogenetic analyses indicating distinct baseline distributions between African and non-African populations. The rapid decay of phylogenetic autocorrelation at short evolutionary distances further supports a dominant role for demographic processes such as drift and population history.

This pattern is consistent with the unusually high mutation rate of the *AMY1* locus, driven primarily by non-homologous allelic recombination events that add or remove copies in discrete units (Groot et al., 1990; Yilmaz et al., 2024). Loss of these mutational units may be effectively irreversible, generating stable low-copy haplotypes and long-lasting population-level differences. Such dynamics could explain the persistence of low average *AMY1* CN in populations such as Siberians and Mayans (Inchley et al., 2016; Bolognini et al., 2024; Scheer et al., 2025), independent of recent dietary practices.

Patterns of decreasing *AMY1* CN diversity with increasing distance from Africa, although preliminary, are also consistent with neutral evolution under serial bottlenecks. Similarly, in ancient European data, correlations between *AMY1* CN, ancestry components, and time suggest that post-Neolithic increases may reflect demographic turnover rather than direct selection on diet alone, paralleling recent findings for lactase persistence in South Asia (Kerdoncuff et al., 2025).

### Implications

Together, our results indicate that *AMY1* CN variation does not represent a universal or straightforward genetic signature of agricultural adaptation. Instead, it reflects a complex interplay of demographic history, mutation dynamics, and potentially diverse selective pressures. More broadly, this study highlights the importance of ancestry-aware analyses and comprehensive geographic sampling when interpreting putative adaptive signals in the human genome, particularly for loci with extreme structural variability.

## Conclusion

Our results show that agriculture is not a universal driver of *AMY1* CN diversity and that previously reported associations likely reflect limited sampling and incomplete control for shared ancestry rather than direct selection on dietary practices. Instead, demographic history, including the dispersal of modern humans out of Africa, subsequent gene flow, and population turnover in different regions, emerges as a more consistent explanation for global patterns of variation at this locus.

By re-evaluating a canonical example of human dietary adaptation using ancestry-aware analyses and expanded geographic sampling, this study revises a long-standing paradigm in human evolutionary genomics. More broadly, it underscores the need to disentangle demography from selection when interpreting putative adaptive signals, particularly at structurally complex loci, and highlights the central role of African diversity in testing evolutionary hypotheses.

## Methods

### Data generation and annotation

*AMY1* copy-number (CN) estimates analyzed in this study were generated using two complementary approaches and are therefore analyzed separately. The first dataset (‘**ddPCR dataset**’) consists of 459 droplet digital PCR (ddPCR) estimates, including 318 Sub-Saharan African individuals reported for the first time in this study and 141 individuals from the Human Genome Diversity Project (HGDP) panel (72 Africans and 69 non-Africans) (Figures 1A and S1; Table S1.1). Two independent single-copy reference loci were used to estimate *AMY1* CN, and estimates were validated using independent data available for 138 HGDP individuals (Bolognini et al., 2024) (Figure S6). Detailed descriptions of the ddPCR protocol, quality control, and filtering procedures are provided in the Supplementary Methods. Biological samples from the HGDP individuals were provided by the CEPH Biobank, Paris, France (BIORESOURCES), while all other samples were independently collected following informed consent and ethical approvals obtained by collaborators in the respective African countries. Ethics reference numbers and additional details are provided in the Supplementary Methods.

The second dataset (‘**read-depth dataset**’) was obtained from Bolognini et al. (2024) and includes 1,307 individuals from the HGDP, the Simons Genome Diversity Project (SGDP), and the 1000 Genomes Project (KGP) (Figures 1B and S2; Table S1.2).

Whole-genome sequencing data for individuals in the read-depth dataset was retrieved from the International Genome Sample Resource, while genotypes from the remaining individuals of the ddPCR dataset was obtained from previous publications (Schlebusch et al., 2012; Vicente et al., 2019; Fortes-Lima et al., 2022, 2024), except for Ethiopian, Bassar (Togo), and Mbo (Cameroon) individuals, whose genomic data are reported here for the first time.

In addition, *AMY1* CN estimates for 533 ancient European individuals were obtained from Bolognini et al. (2025), with corresponding genomic data from the Allen Ancient DNA Resource (Mallick et al., 2024). This dataset (‘**ancient Europeans**’) (Figure S3, Table S1.3) was used exclusively for exploratory temporal analyses and was not included in the main modeling framework.

All individuals from the ddPCR and read-depth datasets were assigned to populations based on self-reported ethnicity or KGP, HGDP and SGDP metadata. Subsistence information was compiled from the D-PLACE database (Kirby et al., 2016), and each population was annotated with its dominant subsistence category and five quantitative estimates of dependence on gathering, hunting, fishing, pastoralism, and agriculture. Where detailed information was unavailable, populations were assigned only a dominant subsistence category. As a result, five African ddPCR populations lack full quantitative subsistence annotations. Population descriptions and annotation levels are summarized in Tables S2.1 and S2.2.

### Modeling *AMY1* copy number variation

To assess the relative contributions of shared ancestry and subsistence strategy to *AMY1* CN variation, we applied two phylogenetically-informed regression frameworks in R (v4.4.2), that explicitly account for non-independence among observations due to population structure. Similar approaches have been used to study cold adaptation in humans (Key et al., 2018).

Bayesian models were implemented using the **brms** package (Bürkner, 2017), which allows the incorporation of individual-level phylogenetic autocorrelation structures through a variance-covariance (VCV) matrix derived from a phylogenetic tree. Complementarily, frequentist models were implemented using the **glmmTMB** package (Brooks et al., 2017), which cannot incorporate correlation matrices directly, but instead can include fixed effects describing genetic ancestry in the form of principal components (PCs). In our modeling setup, the null model included genetic ancestry as either a random (brms) or fixed (glmmTMB) effect, while all other models additionally included subsistence predictors, specified as binary contrasts (agriculturalist vs. non-agriculturalist), multi-level subsistence categories, or continuous measures of subsistence dependence. A summary of all model specifications is provided in Table 1, and detailed descriptions on the tests prior to modeling are provided in Supplementary Methods.

Model assumptions and fits were evaluated following Mundry (2014). Model performance in the frequentist framework was assessed using likelihood ratio tests and 95% confidence set model comparisons (Symonds & Moussalli, 2011). For Bayesian models, predictor effects were evaluated using 95% credible intervals and regions of practical equivalence implemented in **bayestestR** (Makowski et al., 2019).

### Temporal analyses in West Eurasia

To explore temporal changes in *AMY1* CN in Europe, ancestry proportions were estimated for 533 ancient individuals using supervised ADMIXTURE analyses (Alexander et al., 2009), specifying Western hunter- gatherers, Yamnaya pastoralists, and Anatolian farmers as source populations. Associations between ancestry proportions, sample age, and *AMY1* CN were evaluated by regression in glmmTMB. Detailed descriptions of these tests are provided in the Supplementary Methods.

### Out-of-Africa dispersal and copy-number diversity

To investigate whether the demographic expansion out of Africa shaped *AMY1* CN diversity, we calculated five summary statistics of CN diversity for non-Sub-Saharan populations in the read-depth dataset and correlated them with geographic distance from East Africa. Only populations with at least five individuals were included, and diversity estimates of all groups were based on downsampled replicates to control for unequal sample sizes. Detailed descriptions of these tests are provided in the Supplementary Methods.

### Phylogenetic signal analyses

Phylogenetic signal in *AMY1* CN was assessed independently of the modeling framework using the **phylosignal** package (Keck et al., 2016). Analyses included global measures of phylogenetic signal, phylogenetic correlograms, and local indicators of phylogenetic association. These complementary approaches quantify whether trait similarity among individuals departs from random expectations and identify phylogenetic clustering or outlier values across the human phylogeny. Detailed descriptions of these tests provided in the Supplementary Methods.

## Acknowledgements

We thank all sample donors from individual projects across Africa and from the HGDP panel. We are also grateful to the CEPH Biobank, Paris, France (BIORESOURCES) at Fondation Jean Dausset-CEPH for the maintenance of HGDP cell lines and their DNA distribution. We gratefully acknowledge Jennifer Meadows and Hanna Davies for support on the ddPCR method, as well as Hana Merchant, François Keck, Miguel Redondo, David Orme and Will Pearse for guidance on statistical analyses.

The authors would like to acknowledge support of the National Genomics Infrastructure (NGI)/Uppsala Genome Center and UPPMAX for aiding in massive parallel sequencing and computational infrastructure. Work performed at NGI/Uppsala Genome Center has been funded by RFI/VR and Science for Life Laboratory, Sweden. The computation and data handling were enabled by resources provided by the Swedish National Infrastructure for Computing (SNIC) at UPPMAX partially funded by the Swedish Research Council (grant no. 2018-05973). This project was supported by funding to C.S. from the European Research Council (ERC) under the European Union’s Horizon 2020 research and innovation program (grant no. 759933), and the Knut and Alice Wallenberg foundation. C.B. received funding from the European Union’s Horizon 2020 research and innovation program under the Marie Skłodowska-Curie Fellowship Programme (grant no. 839643). Sample collection in the DRC was funded by the ERC-CoG awarded to K.B. for the BantuFirst project (grant no. 724275). A.S.N also received funds from the Swedish Phytogeographical Society (2025) and Helge Ax:son Johnsons Stiftelse (grant no. F25-0211) for the international dissemination of this work and the purchase of computational equipment.

## Declaration of generative AI and AI-assisted technologies in the manuscript preparation process

During the preparation of this work the authors used ChatGPT in order to do language and grammar checks. After using this service, the authors reviewed and edited the content as needed and takes full responsibility for the content of the published article.

## Data and code availability

*AMY1* copy number estimates for all individuals included in this study are provided in Supplementary Materials, including estimates generated in this study (Table S1.1) and in Bolognini et al. (2024) (Table S1.2). Subsistence annotation for all populations is also provided (Tables S2.1 and S2.2). SNP array genotype data of modern-day African populations generated in this project (Arbore, Bassar, Gurage, Hamar and Karo, Mbo, Ongota, and Sheko) is available through the European Genome-Phenome Archive (EGA) data repository subject to Data Access agreements. The remaining samples from the ddPCR dataset can be accessed following the instructions in previous publications (Schlebusch et al., 2012; Vicente et al., 2019; Fortes-Lima et al., 2022; Fortes-Lima et al., 2024). Genotyping data for individuals from the Thousand Genomes Project, the Human Genome Diversity Panel and the Simons Genome Diversity Panel can be found in the International Genome Sample Resource database (https://www.internationalgenome.org/). Descriptions of all analyses and corresponding bash and R code have been included in Supplementary Methods. Both Supplementary Methods and Supplementary Materials will be uploaded to GitHub upon acceptance (https://github.com/Schlebusch-lab/AMY1-CN).

---

**Extended data S1.**
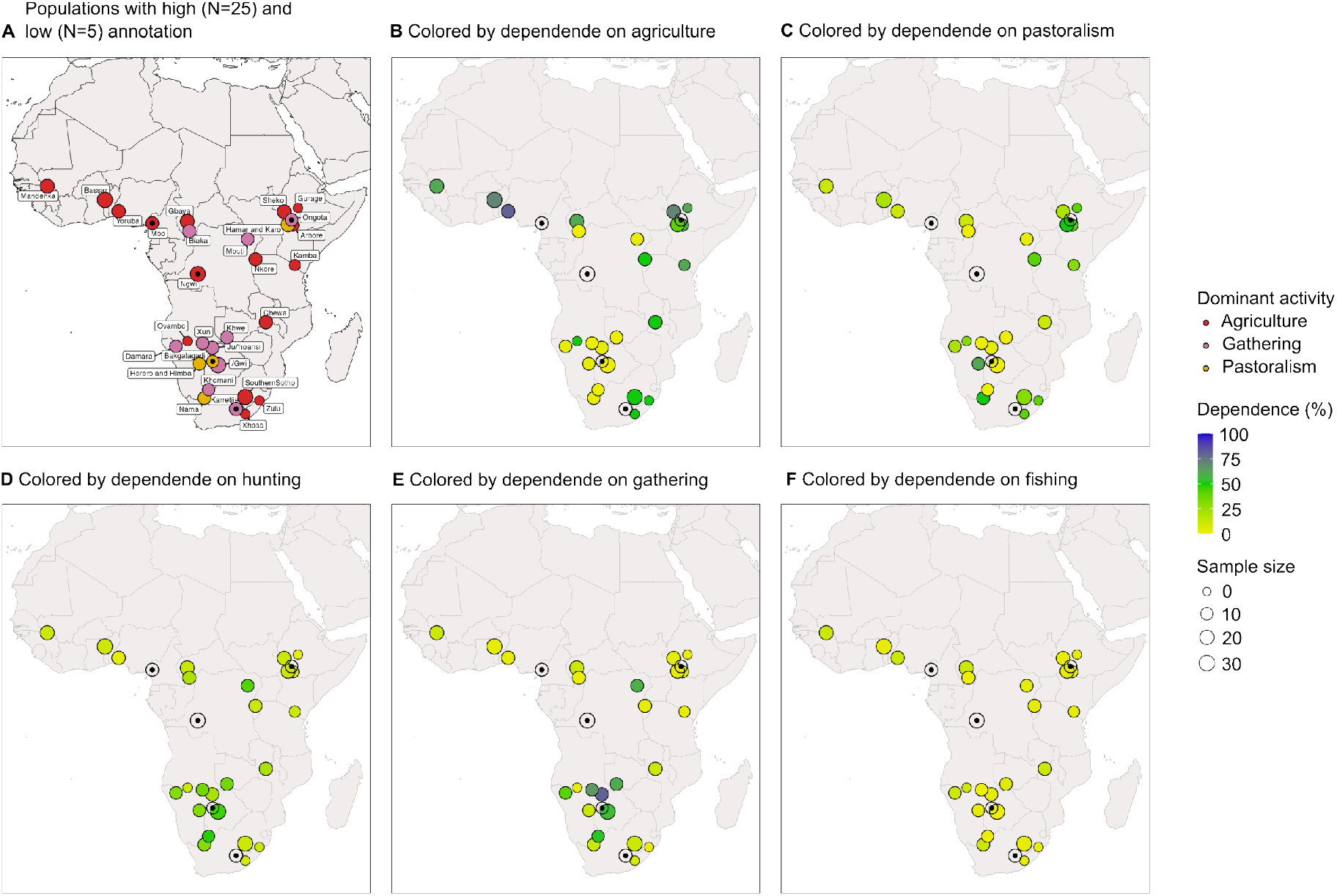
Description of the populations in the African **ddPCR data set**. Only the first panel (**A**) shows the low annotation populations (Bakgalagadi, Karretje, Mbo, Ngwi, Ongota), which are marked with a black dot. The remaining panels show the high level annotation populations colored by their dependence on agriculture (**B**), animal husbandry or pastoralism (**C**), hunting (**D**), gathering (**E**) and fishing (**F**), with circle sizes corresponding to population sample size.

**Extended data S2.**
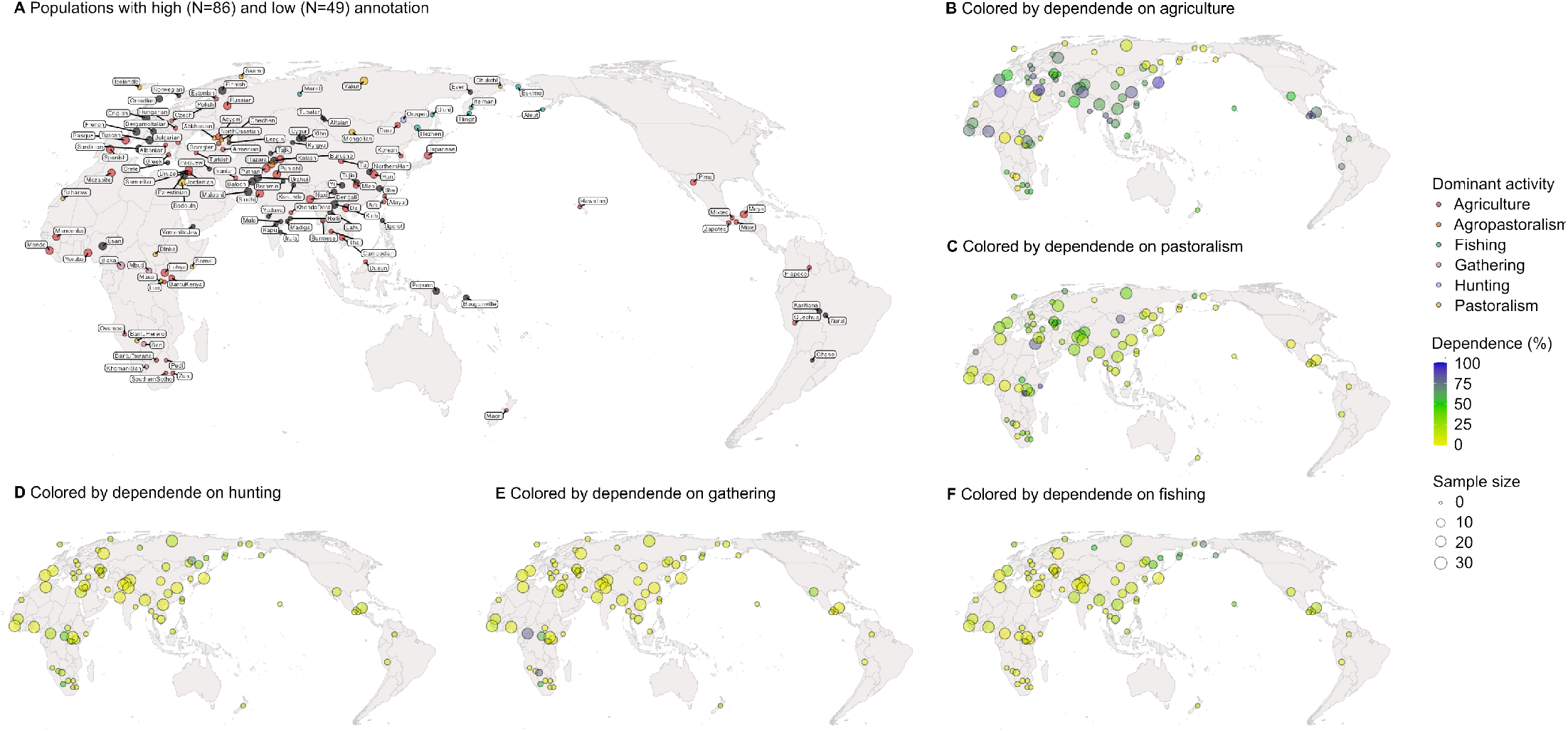
Description of the populations of the **read-depth data set**. Only the first panel (**A**) shows the low (N=86) and high (N=49) annotation populations. The remaining panels show only the highly annotated populations colored by their dependence on agriculture (**B**), animal husbandry or pastoralism (**C**), hunting (**D**), gathering (**E**) and fishing (**F**), with circle sizes corresponding to population sample size.

**Extended data S3.**
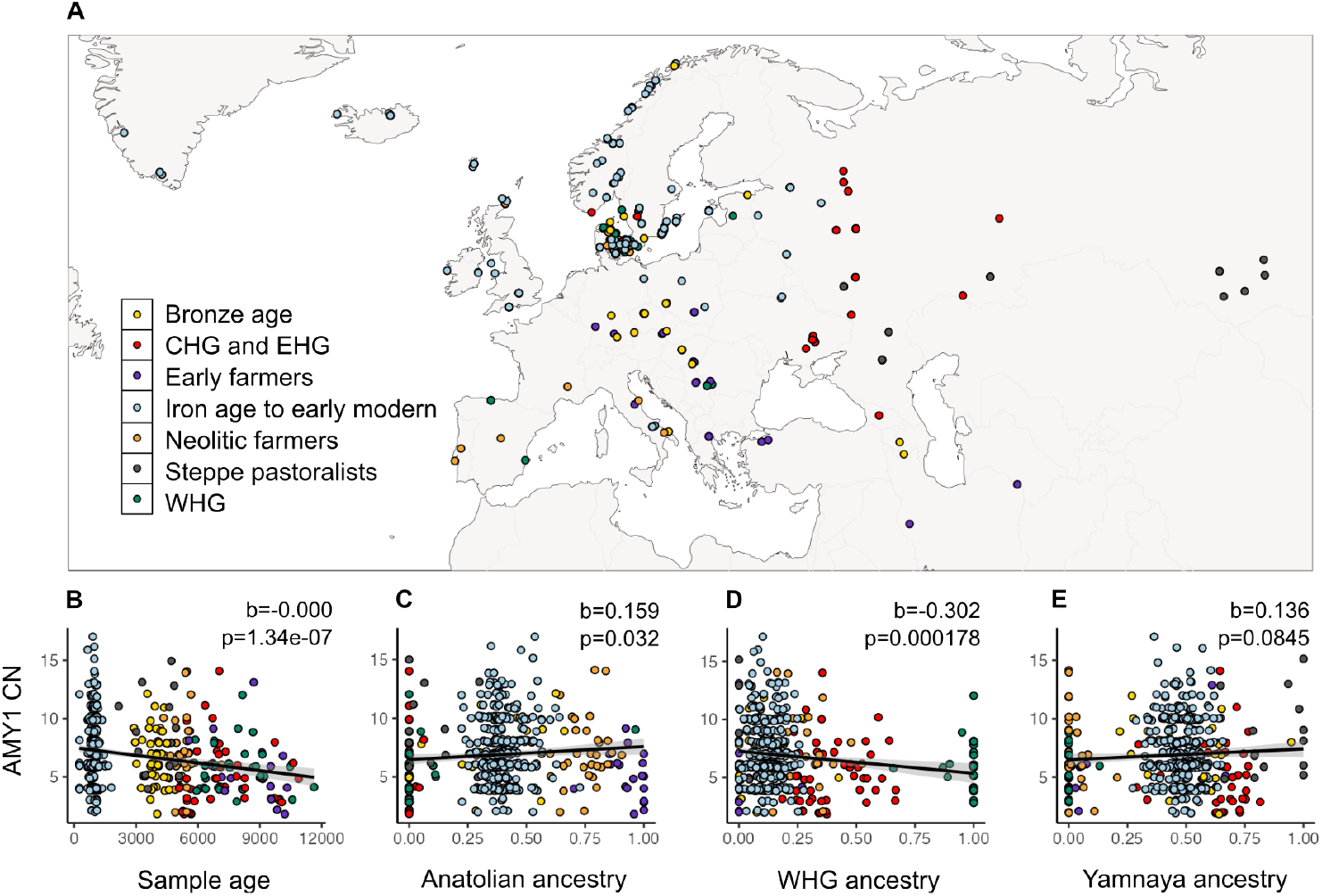
Geographical location of 533 ancient Europeans reported in Bolognini et al. (2024) (**A**) and correlation between their *AMY1* CN and age (∼12,000 to ∼500 years ago) (**B**), as well as their ancestry proportions from a supervised admixture with three sources: Anatolian farmers (**C**), Western hunter-gatherers (WHG) (**D**) and Yamnaya pastoralists (**E**). Regression coefficients (b) and associated p-values (p) correspond to generalized linear mixed models. Black trend lines correspond to isolated linear models between the plotted variables.

**Extended data S4.**
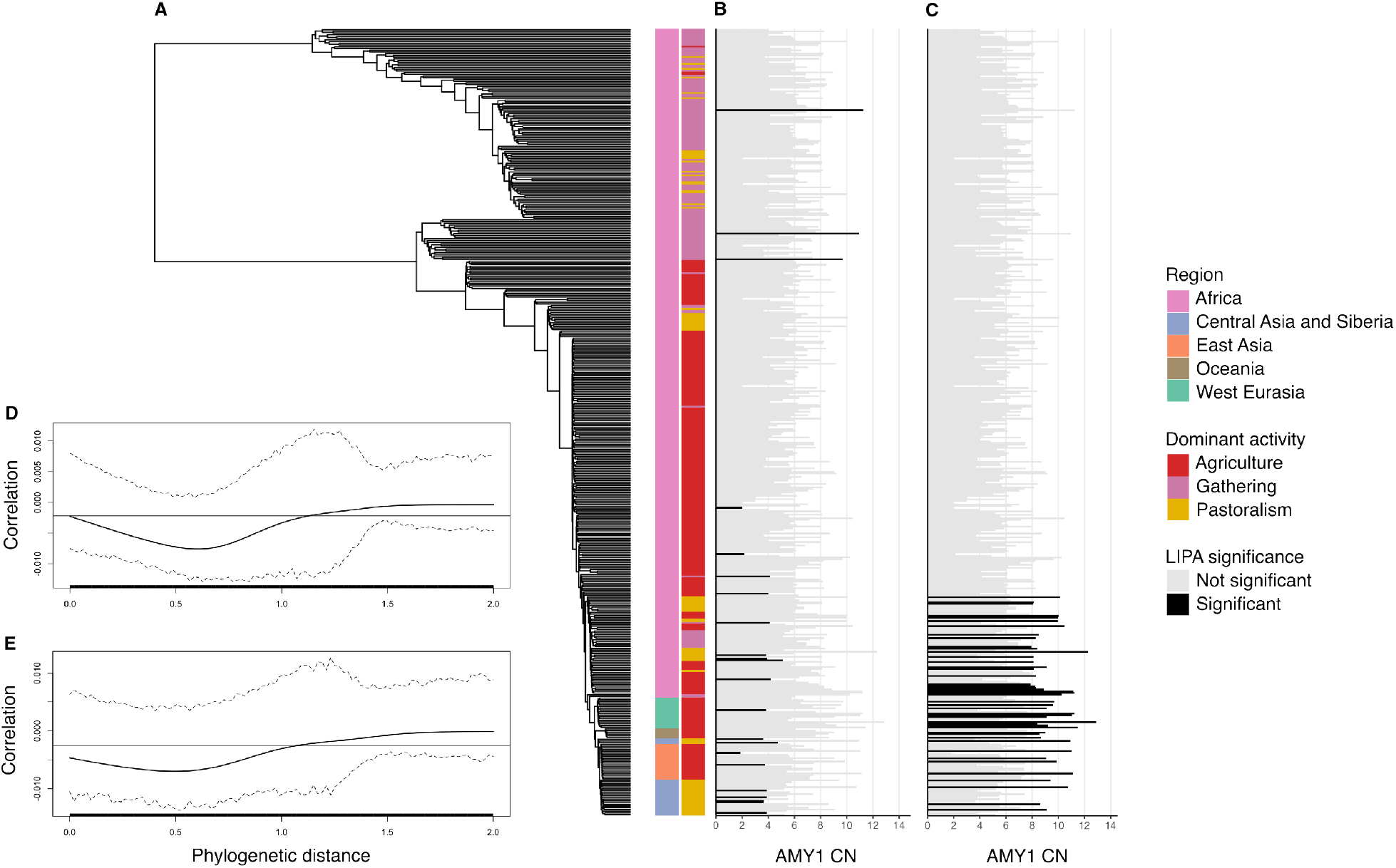
Phylogenetic tree of the global **ddPCR data set** (n=459, N=35) (**A**), with results from the local indicators of phylogenetic association (LIPA) tests (**B, C**) and phylogenetic correlograms (**D, E**). Each terminal branch in the phylogenetic tree (**A**) corresponds to an individual in the **ddPCR data set** and the two associated colored plots correspond to the geographical region (left) and dominant activity (right) of the population to which each individual belongs. Individual *AMY1* CN and results of the LIPA test for the ‘less’ and ‘greater’ hypotheses are shown in the second (**B**) and third (**C**) panels, respectively. Each bar is colored by the significance of its LIPA test (alpha=0.05). The remaining panels (**D, E**) show results of the phylogenetic correlogram tests using the complete data set (n=459, N=35) (**D**) and all African populations alone (n=390, N=30) (**E**).

**Extended data S5.**
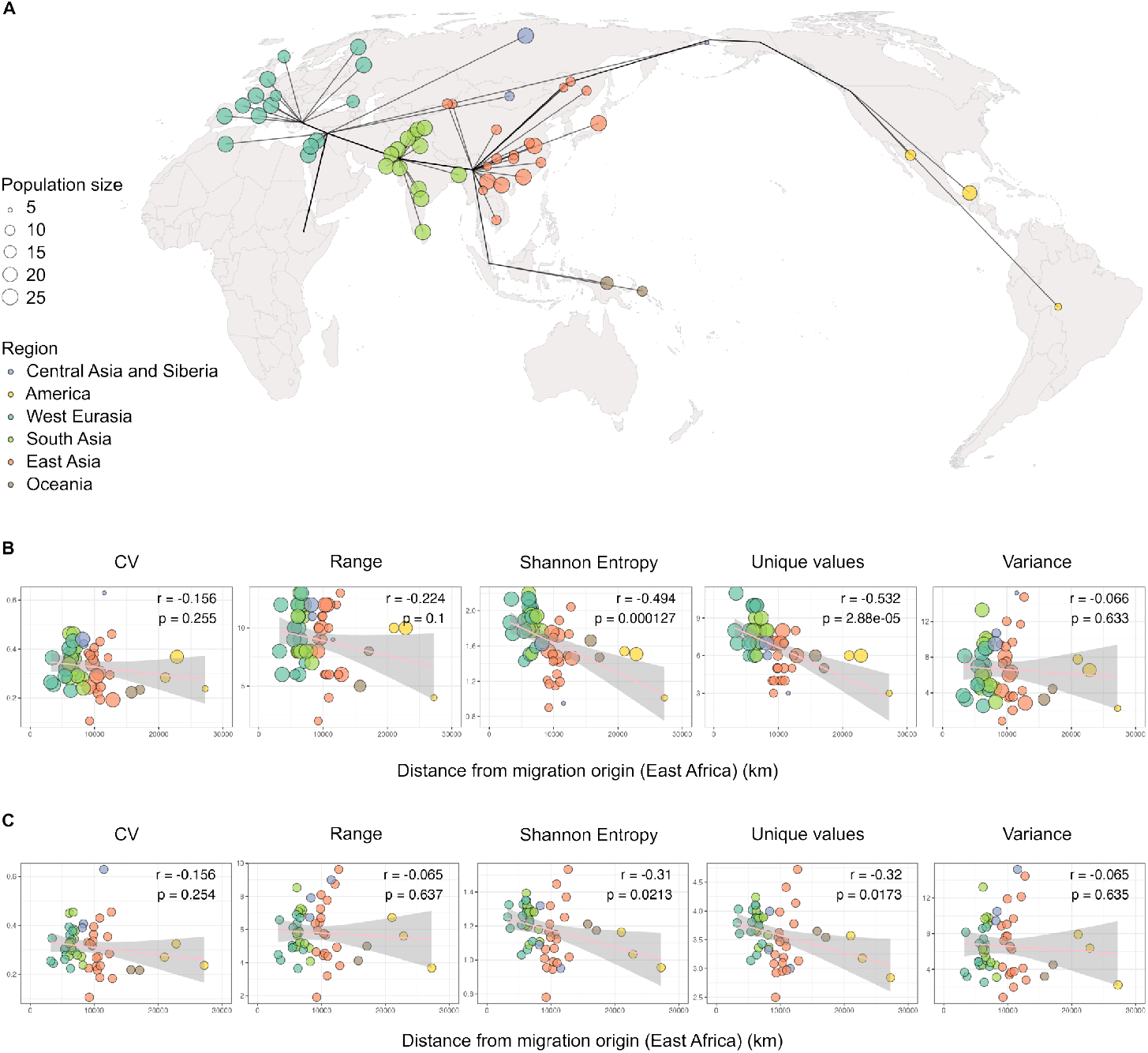
Correlation between distance to East Africa and several measures of diversity in *AMY1* CN. The first panel (**A**) illustrates the geographical distances between each population and East Africa. Populations are colored by region and their sample size is represented by the size of the colored circles. Plots in the panels below (**B, C**) show the relationship between geographical distance to East Africa and five measures of diversity (CV: coefficient of variation; range; Shannon entropy; number of unique values; Var: variance) for the original populations (**B**) and for 500 replicates of populations downsampled to 5 individuals each (**C**). In both approaches, Shannon entropy and number of unique values are negatively correlated with distance from East Africa.

**Extended data S6.**
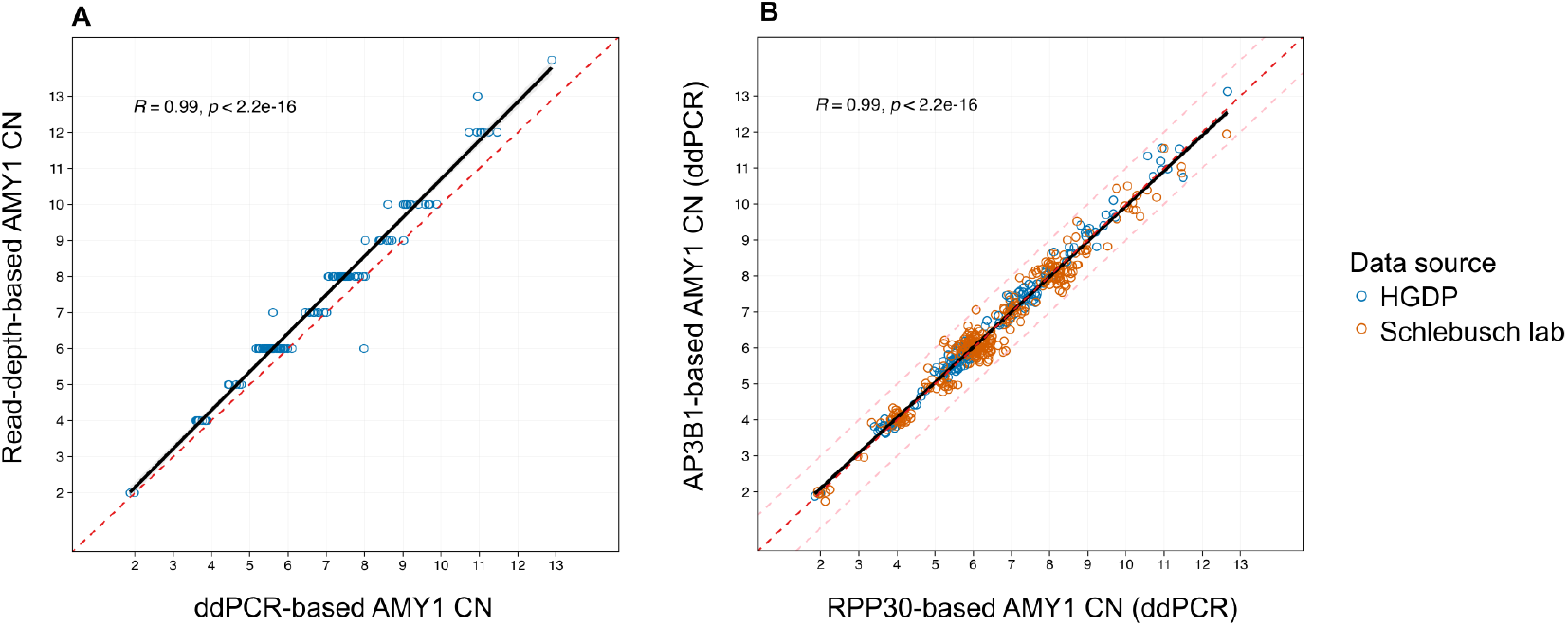
Correlation between ddPCR and read-depth *AMY1* CN estimates for 138 individuals of the HGDP panel (**A**) and between ddPCR estimates generated with *AP3B1* and *RPP30* reference loci (**B**). Correlation (R) and p-values (p) refer to a Pearson’s correlation test and the black trend line represents an isolated linear model between the two plotted variables. The red dashed line marks a 1:1 relationship, and the pink dashed lines in the second panel (**B**) delimit the space where the absolute difference between estimates obtained with each reference locus is less than one.

