## Supplementary Methods for "Rethinking human *AMY1* copy number evolution in light of demographic history"

2026-01-20

#### 1 Sample origin and ethical permits

##### 1.1 HGDP samples

The 'Centre d'Etude du Polymorphisme Humain' Biobank, Paris, France (BIORESOURCES), provided extracted DNA from lymphoblastoid cell lines from all individuals in the Human Genome Diversity Panel (HGDP) used in this study. Purchase was approved on January 11th, 2021, by the biobank operational manager.

##### 1.2 Non-HGDP samples

Non-HGDP biological samples for this study were collected by our collaborators, who obtained the original ethical permission for the sampling in African countries. **Table 1** contains the ethics reference numbers, and local and Swedish permission details associated with these samples. All participants donated saliva samples with informed consent.

**Table 1.** Sample origin and corresponding ethical permits.

| Collection name | Country | Swedish clearance | Local clearance | Local clearance institution |
|---|---|---|---|---|
| DRC samples | DRC | Dnr 2019-05244 and Dnr 2021-01448 | 091/CAB/MIN/-CA/PKB/2018 and CAB/MIN/CAP/-JJM/JLM/051/22 | Minister of Arts and Culture of the DRC |
| Ethiopian samples | Ethiopia | Dnr 2021-01448 | 310/169/2018 | The Federal Democratic Republic of Ethiopia Ministry of Science and Technology |

| Collection name | Country | Swedish clearance | Local clearance | Local clearance institution |
| --- | --- | --- | --- | --- |
| Khoe-San and Bantu samples from Africa | Zimbabwe, South Africa, Namibia, Uganda, Zambia, Zanzibar | Dnr 2021-01448 | M180654, M180655, M180656 | Human Reearch Ethics Committee from Witwatersrand University |
| Khoe-San and Bantu samples from Botswana | Botswana | Dnr 2019-00115 | Permission obtained from the Botswana government in accordance with national regulations in place at the time of sampling | Botswana governmental office |
| South African samples | South Africa | Dnr 2021-01448 | EC160429-024 (Health Ethics reference 259/2016) | Faculty of Natural and Agricultural Sciences University of Pretoria |
| Cameroon samples | Cameroon | Dnr 2021-01448 | CBI/397/ERCC/-CAMBIN | Ethics Review and Consultancy Committee (ERCC), Cameroon Bioethics Initiative (CAMBIN) |
| Togo samples | Togo | Dnr 2021-04382 | N097/MESR/SG/-DRST/19 | Ministry of higher education and research |

### 2 *AMY1* copy number estimation by droplet-digital PCR

#### 2.1 Droplet-digital PCR

Droplet-digital PCR (ddPCR) provides absolute quantification of a genomic target by partitioning the DNA sample into ~20,000 droplets and scoring each individual droplet for fluorescence signals after PCR amplification. Fluorescence is only emitted by TaqMan probes when target-specific primers anneal to the target region and are elongated during the PCR. Droplets that emit fluorescence signals are read as positive, while droplets that do not are read as negative. Two fluorescence assays must be included in each ddPCR reaction, i.e. one targeting the genomic region of interest, the other targeting a reference locus of known copy number (CN). The fraction of positive droplets for each of the assays is fitted to a Poisson distribution to finally determine the absolute initial CN of the target DNA.

#### 2.2 Copy number estimation

In this study, we estimated *AMY1* CN of each individual using not one but two single-copy reference loci (*RPP30* and *AP3B1*) in separate runs of the BioRad Laboratories assay (for *AMY1*: dHsaCP1000594; *RPP30*: dHsaCP1000485; *AP3B1*: dHsaCP1000001). DNA was digested in 10 U:100 ng enzyme:DNA proportions with HaeIII restriction enzyme (10 U/$\mu$l, NewEngland Biolabs, Cat#:R0108L) in 10X buffer rCutSmart (NewEngland Biolabs, Cat#:B6004S) for 1 h at 37°C. A 22 $\mu$l mixture of 2×ddPCR Supermix (Bio-Rad, Cat#:186–3024), forward and reverse primers for target and reference assay (final concentrations of 900 nM each), probes for both assays (final concentrations of 250 nM each) and 15 ng of digested DNA was emulsified with Bio-Rad Droplet Generator Oil (Bio-Rad, Cat#:186-4110) in ddPCR™ 96-well plates

(Bio-Rad, Cat#:12001925) in a Bio-Rad QX200 Automated Droplet Generator (Bio-Rad, Cat#:186–4101) according to the manufacturer's instructions. Before and after droplet generation, the plates were heat-sealed with sealing foil sheets (BioRad, Cat#:181-4040) using the Bio-Rad PX1 Plate Sealer (Bio-Rad, Cat#:181–4000) at 180 °C for 5 s. PCR was performed with the following cycling parameters: 10 min at 95 °C (1 cycle), 30 s denaturation at 94 °C and 1 min annealing and extension at 60 °C (40 cycles), 10 min at 98 °C and a hold at 4 °C. All steps had a ramp rate of 2 °C/s.

Two negative controls were included in each plate, and several samples were run across different plates. Droplets were analyzed using a Bio-Rad QX200 Droplet Reader (Bio-Rad, Cat#:186–4003). Fluorescent data from each well was analyzed with `QuantaSoft` (version 1.7.4) based on a Poisson distribution. Droplet generation and copy number estimation was performed at the Department of Immunology, Genetics and Pathology of Uppsala Biomedical Centre (BMC, Uppsala University).

##### 2.3 Filtering and validation of estimates

We successfully generated a total of 1200 ddPCR estimates for 591 individuals. Of these estimates, 589 were based on the *RPP30* reference locus and 611 on *AP3B1*. Of the 591 total individuals, 43 didn't have estimates based on both reference loci and were therefore filtered out. Estimates from the remaining 548 individuals were used to plot the correlation between *RPP30*-based and *AP3B1*-based AMY1 CN, which showed very high accordance (P corr=0.97, p-value<0.001) and the presence of a few discrepancies. These discrepancies were flagged if the absolute difference between *RPP30*-based and *AP3B1*-based *AMY1* CN estimates was bigger than one:

$$|\text{RPP30-based AMY1 CN}–\text{AP3B1-based AMY1 CN}| > 1$$

Discrepant estimates for individuals with just one estimate per reference locus were discarded, due to not being able to identify which of the two estimates was more accurately describing the true *AMY1* CN of that individual. This was the case for only nine individuals. Individuals with more than one estimate per reference locus were retained if we could easily deduce the true *AMY1* CN estimate (e.g. by identifying a single *AP3B1*-based or *RPP30*-based estimate that was different from all other estimates for that individual and therefore more likely to be incorrect). After this quality control, the remaining number of individuals with a high confidence *AMY1* CN estimate was 545. Due to kinship filtering and the removal of individuals with mixed ancestry, unclear group label or lack of whole-genome genotyping data, the total number of individuals we proceeded with was brought down to 459.

Correlation plots and tests were repeated with the filtered data set and, as expected, showed improvement in the relationship between *AP3B1*-based and *RPP30*-based estimates (P corr=0.99, p-value<0.001). Here, we noted that *AMY1* CN estimates from HGDP samples tend to receive values more distant from integers than non-HGDP samples, which we argue could be due to CN losses and gains over the maintenance of HGDP cell lines. Additionally, because the **ddPCR** and **read-depth data sets** overlap for 138 samples from the HGDP panel, we validated the ddPCR *AMY1* CN estimates by testing and plotting the correlation between estimates originating from the two methods (P corr=0.99, p-value<0.001). We noted that ddPCR-based estimates tend to be somewhat lower than read-depth-based estimates for the same individuals, especially at higher values of *AMY1* CN, again pointing at HGDP cell-line instability at least for this locus.

After the filtering steps, we averaged the *RPP30*- and *AP3B1*-based *AMY1* CN for all 459 individuals kept in the data set and used this value in all subsequent tests. All ddPCR estimates in the final data set are available in **Supplementary Materials** (**Table S1.1**), and correlation plots for the estimates validation are shown in **Figure S6**.

#### 3 Collection of genotyping data

Genome-wide data of individuals belonging to the **Thousand Genomes Project** (KGP), the **Human Genome Diversity Project** (HGDP) and the **Simons Genome Diversity Project** (SGDP) was obtained from the **International Genome Sample Resource** (IGSR) database:

- https://www.internationalgenome.org/

The remaining samples from the **ddPCR data set** can be accessed following the instructions in previous publications (Schlebusch et al., 2012; Vicente et al., 2019; Fortes-Lima et al., 2022; Fortes-Lima et al., 2024), except for all 39 Ethiopian, the 28 Bassar (Togo) and the 13 Mbo (Cameroon) individuals, whose genotyping data is published for the first time under this study. DNA samples corresponding to the latter individuals were genotyped at the SNP&SEQ Technology Platform, NGI/SciLifeLab Genomics (Sweden), on the Illumina Infinium H3Africa Consortium genotyping array.

- (EGA ID will be provided upon acceptance)

Genotyping data for the 533 ancient European samples was obtained from the **Allen Ancient DNA Resource** (AADR) (Mallick et al., 2024). Database version 62.0 typed on variants of the 1240k panel was used.

- https://dataverse.harvard.edu/dataset.xhtml?persistentId=doi:10.7910/DVN/FFIDCW

### 4 Filtering of data sets

#### 4.1 Genetic relatedness

Kinship analyses were run on both data sets with `KING` (version 2.3.2) (Manichaikul et al., 2010) to identify relatives up to the second-degree (corresponding to a kinship coefficient of 0.0884). For each pair of related individuals, one of them was randomly selected and removed. SNP variants across the whole genome were used in this test.

```
# PLINK file set comprised of .bed, .bim and .fam files
DB=ddPCR_data_set

# Run KING
/king -b ${DB}.bed --kinship --prefix KING/${DB} > KING/${DB}_king_info.txt

# Sort individuals based on their kinship coefficient
sort -k9 -r -h KING/${DB}.kin > KING/${DB}_kinship_list.txt

REL_N=20 # Ex. amount of related pairs (20)

# Identify (unique) related individuals
head -n ${REL_N} KING/${DB}_kinship_list.txt | \
    cut -f3 | sort | uniq > related_list.txt

# Get their full names from the .fam file
cat related_list.txt | while read line; \
    do grep ${line}' ' ${DB}.fam >> related_list_full_names.txt; \
    done

# Remove them from dataset
plink --bfile ${DB} --remove related_list_full_names.txt --make-bed \
    --out ${DB}_unrelated
```

#### 4.2 Disparate population sizes

Additionally, for the **read-depth data set**, populations with more than 25 individuals (mostly those originating from the KGP) were randomly downsampled to 25 to reduce the disparate range of population sizes, once related individuals had been removed. The final population sizes are summarized in the **Table 2**.

**Table 2**. Summary of population sizes.

| Data set | Nr samples | Nr pops | Avg. pop size | Median pop size | Min. and max. pop size |
|---|---|---|---|---|---|
| ddPCR | 459 | 35 | 13.11 | 13 | (1,28) |
| read-depth | 1307 | 135 | 9.68 | 3 | (1,25) |

### 5 Annotation of data sets

Each individual included in either of the data sets was assigned to a population, following the KGP, HGDP and SGDP descriptions for the **read-depth data set** or the self-reported ethnicity of the remaining individuals in the **ddPCR data set**. The geographical location of these populations was retrieved from either the KGP, HGDP and SGDP metadata descriptions, the location where DNA sampling took place or the location where the majority of an ethnicity currently lives.

- KGP: https://www.internationalgenome.org/data-portal/data-collection/phase-3
- HGDP: https://www.internationalgenome.org/data-portal/data-collection/hgdp
- SGDP: https://www.internationalgenome.org/data-portal/data-collection/sgdp

Whenever possible, detailed information on the subsistence of each population was obtained from the Database of Places, Language, Culture and Environment (D-PLACE) (Kirby et al., 2016) (https://d-place.org/), which provides comprehensive descriptions of many variables of anthropological interest. The variables used in this study are described in **Table 3**. Mathematical transformations of the dependency variables are explained in **Section 6**.

**Table 3**. Variable specifications.

| Variable name | Variable ID and descriptions (D-PLACE) |
|---|---|
| **Dominant activity** | [B004] Subsistence activity that provides the majority of a group's nutritional intake (hunting, gathering, or fishing).<br>[EA042] Dominant mode of subsistence (note: not in original EA; derived from other variables on subsistence and type of agriculture).<br>[SCCS246] Identical to EA042. |
| **Dependency on agriculture** | [EA005] Dependence on agriculture, relative to other subsistence activities.<br>[SCCS207] Identical to EA005. |
| **Dependency on pastoralism** | [EA004] Dependence on animal husbandry, relative to other subsistence activities.<br>[SCCS206] Identical to EA004. |
| **Dependency on hunting** | [B002] Group's dependence upon hunting of terrestrial animals, relative to other subsistence activities.<br>[EA002] Dependence on hunting, including trapping and fowling, relative to other subsistence activities.<br>[SCCS204] Identical to EA002. |

| Variable name | Variable ID and descriptions (D-PLACE) |
|---|---|
| **Dependency on gathering** | [B001] Group's dependence upon gathering of terrestrial plants, relative to other subsistence activities.<br>[EA001] Dependence on the gathering of wild plants and small land fauna, relative to other subsistence activities.<br>[SCCS203] Identical to EA001. |
| **Dependency on fishing** | [B003] Group's dependence upon fishing of aquatic organisms, relative to other subsistence activities.<br>[EA003] Dependence on fishing, including shell fishing and the pursuit of large aquatic animals, relative to other subsistence activities.<br>[SCCS205] Identical to EA003. |

### 6 Modeling *AMY1* CN based on subsistence and demography

Comparing *AMY1* CN among groups of different subsistence strategies without taking into account the genetic background of the included individuals can falsely attribute the detected difference to the subsistence strategy of the compared groups. In other words, differences due to a variable that is unaccounted for (e.g. genetic relatedness between individuals) will be wrongly attributed to the included predictor variable (e.g. subsistence strategy). Accounting for the lack of independence between observations can be done in numerous ways. In genome-wide association studies, shared ancestry is usually considered through principal component analysis (PCA), multidimensional scaling or genetic relatedness matrices (Uffelman et al., 2021; Marees et al., 2018).

In the main text, we briefly describe the implementation of two different modelling methods that can deal with non-independent observations by either (i) incorporating the expected pattern of auto-correlation between individual measures in the form of a variance-covariance (VCV) matrix, or (ii) controlling for genetic similarity between observations with the addition of fixed effects that describe ancestry components resulting from PCA. In this section, we first describe the model fits, their evaluation and comparison, and later report how the data was formatted for appropriate use in the implementation of each of these methods. Two examples where a methodology conceptually similar to ours has been used to investigate *allele frequency* variation in the context of adaptation to climate in humans are Key et al., 2018 and Grover-Thomas et al., 2025.

#### 6.1 Model fits

##### 6.1.1 Response variable

In both `brms` (Brükner, 2017) and `glmmTMB` (Brooks et al., 2017) modeling approaches, the response variable was *AMY1* CN as obtained by the ddPCR (non-integer) or the read-depth methods (integer). Due to its integer, count-like nature, *AMY1* CN from the **read-depth data set** was modeled with either a compois (in `glmmTMB`) or a Poisson (in `brms`) error structure with their default log link functions. On the other hand, continuous-like *AMY1* CN of the **ddPCR data set** was modeled with either the default Gaussian error structure (in `glmmTMB`) or a Student's t-distribution (in `brms`), which behaves similarly to a Gaussian distribution but is more robust to possible outliers.

##### 6.1.2 Predictor variables (fixed and random effects)

The five predictor variables describing a population's dependency on each subsistence strategy (*agriculture*, *pastoralism*, *hunting*, *gathering*, *fishing*) are provided in D-PLACE as a range of values (e.g., 56-65%). For all these variables and populations, we kept only the intermediate value (e.g., 60.5% in the previous case) and transformed it by the isometric log ratio (ILR), a common approach to ensure that the relative proportions of components are appropriately modeled, avoiding issues of spurious correlations inherent in raw compositional data.

- **Zero replacement**: Since compositional data cannot contain zeros, we applied the count zero multiplicative (CZM) replacement method using the `cmultRepl` function from the `zCompositions` package (Palarea-Albaladejo & Martín-Fernández, 2015) when necessary.
- **ILR transformation**: We constructed ILR coordinates by computing log-ratios using orthonormal contrast vectors. This way, each ILR coordinate represents a log-ratio between one component and the geometric mean of the remaining components, i.e., the increase or decrease in that subsistence strategy relative to the others.

```
# Select compositional variables
comp_vars <- ddPCR_data_set[,c("agriculture", "pastoralism", "fishing",
                              "gathering", "hunting")]

# Replace zeros if present
comp_vars_no_zeros <- if (any(comp_vars == 0)) {
  cmultRepl(comp_vars, method = "CZM", output = "p-counts")} else {comp_vars}

# Convert to compositional class
comp <- acomp(comp_vars_no_zeros)

# Number of variables
D <- ncol(comp)

# FUNCTION: Create ILR contrast vectors and compute ILR coordinates (new variables)
create_ilr_vector <- function(i, D) {
  v <- rep(-sqrt(1 / (D * (D - 1))), D)
  v[i] <- sqrt((D - 1) / D)
  return(v)}

log_comp <- log(comp)                                                # Log of variables
log_comp_mat <- matrix(unclass(log_comp), nrow = nrow(log_comp), ncol = D)   # Matrix format
colnames(log_comp_mat) <- colnames(comp)                             # Column names

ilr_coords <- matrix(NA, nrow = nrow(ddPCR_data_set), ncol = D)      # Empty matrix
colnames(ilr_coords) <- paste0(colnames(comp_vars), "_ILR")          # Column names

# Apply function to calculate IRL transformations
for (i in 1:D) {
  v <- create_ilr_vector(i, D)
  ilr_coords[, i] <- log_comp_mat %*% v}

# Append ILR coordinates to original dataset
data <- cbind(ddPCR_data_set, ilr_coords)
```

All other fixed effects were incorporated into the models without major transformations. Principal components (PC) obtained from PCA were Z-scaled to amend possible disparate range values. Both categorical variables (*population identity* and *dominant subsistence strategy*) did not need any processing other than ensuring they were treated as factors.

Population identity was included as a random intercept in all models to account for the non-independence of subsistence strategies within populations.

The individual-level phylogenetic VCV matrix was incorporated into the modeling function `brm` of `brms` package as the covariance structure of the individual-level random effect using the `gr(individual, cov=VCV)` syntax.

No interactions were included among any predictors.

###### 6.1.3 Bayesian models specifications

In all `brms` models, we used four chains and 4,000 iterations, of which the initial 3,000 were warm-up. Due to the complexity of the models, parameters `adapt_delta` and `max_treedepth` were increased to 0.999 and 12, respectively, to improve the efficiency and stability of posterior sampling. Model-specific default priors were assigned to all parameters, except for the standard deviation of the phylogenetic random effect (VCV term), which was assigned a somewhat informative yet conservative prior of `normal(0,0.5)` to reduce model instability.

###### 6.1.4 Models formulation

Below are two examples of the `brms` and `glmmTMB` syntax used in modeling *AMY1* CN in the **ddPCR data set**.

```
# Example of glmmTMB model synthax
model_null <- glmmTMB(AMY1_CN ~ PC1 + PC2 + PC3 + PC4 + (1|population),
                      data=ddPCR_data_set,
                      family=gaussian)

# Example of brms model synthax
model_null <- brm(AMY1_CN ~ (1|gr(sample,cov=VCV)) + (1|population),
                  data = brms_data,
                  data2=list(VCV=VCV),
                  prior=set_prior("normal(0,0.5)", class="sd", group="sample"),
                  family=student(),
                  save_pars=save_pars(all=TRUE),
                  sample_prior=TRUE,
                  chains=4, iter=4000, warmup=3000,
                  control=list(adapt_delta=0.999, max_treedepth=12))
```

##### 6.2 Model assumptions and stability

Our data and models were examined following recommendations in Mundry (2014) to evaluate if formal and informal assumptions of linear regression were met. Before fitting the models, we assessed the number of predictors in relation to sample size, the distribution of quantitative predictors and the frequency of categorical predictors in both data sets, which revealed no problems other than data sets being mostly dominated by agriculturalists, which merely reflects the case of present-day populations across the globe.

###### 6.2.1 `glmmTMB` specifics

Specifically for the `glmmTMB` models, we investigated overdispersion, distribution and homogeneity of the model residuals with the `DHARMa` package (Hartig, 2024). We also inspected possible collinearity by variance inflation factor (VIF) using the `check_collinearity` function of the `performance` package (Lüdecke et al., 2021).

```r
# Models assumptions and stability (glmmTMB)
for (name in model_names) {
  # 'models' is a list of all models run with the same data set
  # (e.g. model_null, model_agriculture, model_pastoralism)
  model <- models[[name]]

  # Coefficients summary
  summary(model)$coefficients

  # Multicollinearity check
  check_collinearity(model)

  # Overdispersion test
  sim <- simulateResiduals(model)
  testDispersion(sim)

  # DHARMa diagnostic plots
  plot(sim)
}
```

##### 6.2.2 brms specifics

For the `brms` models, we assessed effective sample sizes (ESS), R-hat, energy Bayesian fraction of missing Information (E-BFMI), Pareto k values, and model convergence using standard diagnostics of the `brms` and `bayesplot` (Gabry & Mahr, 2025) packages. Results automatically obtained by the `brms` package can be found in **Tables S3.1-S3.11**, for all data sets and models evaluated.

```r
# Models assumptions and stability (brms)
for (name in model_names) {
  model <- models[[name]]

  # Summary, LOO and Pareto K
  summary(model)
  loo(model)

  # Sampler diagnostics
  nuts_params <- rstan::get_sampler_params(model$fit, inc_warmup = FALSE)
  nuts_df <- do.call(rbind, nuts_params)

  # Divergences
  if ("divergent__" %in% colnames(nuts_df)) {
    cat("Divergences:", sum(nuts_df[, "divergent__"]), "\n")}

  # Treedepth
  if ("treedepth__" %in% colnames(nuts_df)) {
    max_td <- attr(model$fit, "stan_args")[[1]]$control$max_treedepth
    td_exceeded <- sum(nuts_df[, "treedepth__"] >= max_td)
    cat("Max treedepth exceeded:", td_exceeded, "(", max_td, ")\n")}

  # E-BFMI
  ebfmi <- sapply(nuts_params, function(chain) {
    sum(diff(chain[, "energy__"])^2) / (var(chain[, "energy__"]) * (nrow(chain) - 1))})
  print(ebfmi)
```

```
  # MCMC diagnostics
  mcmc_plot(model, type = "areas", prob = 0.95)
  pairs(model)

  # Multicollinearity
  check_collinearity(model)
}
```

##### 6.3 Model comparisons

To test the performance of different models and consequently evaluate the relevance of the predictor variables that each model includes, we performed comparisons between a null model (which only accounts for genetic ancestry) and a set of nested models (which account for ancestry and additional predictor variables, e.g. subsistence dependencies).

###### 6.3.1 `glmmTMB` specifics

The evaluation of models generated by `glmmTMB`, which uses a frequentist statistical framework, was performed by two methods.

The **95% confidence set** (95CS) approach (Symonds & Mussalli, 2011) identifies the best model(s) among the tested models with 95% probability based on their Akaike's Information Criteria (AIC) score.

```
# Compute AIC values and weights
AIC_values <- sapply(models, AIC)
delta_AIC <- AIC_values - min(AIC_values)
weights <- exp(-0.5 * delta_AIC)
weights <- weights / sum(weights)

# Identify models in the 95% confidence set
sorted_weights <- sort(weights, decreasing = TRUE)
cumulative_weights <- cumsum(sorted_weights)
best_models <- names(sorted_weights[cumulative_weights <= 0.95])
```

The **Likelihood ratio test** (LRT) was performed with `anova` function of the `stats` package between nested models run with the same data.

```
# LRT
anova(null_model, Agr.vs.NonAgr_model)
```

###### 6.3.2 `brms` specifics

These methods are not applicable to models generated under a Bayesian statistical framework, and therefore evaluation of `brms` models was not performed. For all `brms` models, we assessed whether the effect of each predictor was effectively different than zero by (i) the 95% **credible interval** (CI) around its estimate and (ii) the **region of practical equivalence** (ROPE) method, using the `rope` and `equivalence_test` functions of `bayestestR` package (Makowski et al., 2019). ROPE analyses were only deemed relevant when the 95% CI of a given estimate did not include zero.

```r
# ROPE
rope_result <- rope(model)
plot(rope_result)

equivalence_test(model, verbose = FALSE)
```

##### 6.4 Phylogenetic tree

A distance matrix based on the identity-by-state metric was computed for all pairs of individuals in both the **ddPCR** and **read-depth data sets**, using `PLINK` (version 1.90b3n) (Chang et al., 2015) with function `distance`. For this, all variant sites were restricted to chromosome 1 (`--chr 1`), to reduce data size and better represent the phylogenetic history of the amylase locus, and markers with any level of missingness (`--geno 0`) or minor allele frequency below 0.1 (**read-depth data set**) or 0.2 (**ddPCR data set**) (`--maf 0.1`, `--maf 0.2`) were filtered out in order to remove incomplete marker genotypes and variation private to a minority individuals.

```bash
# PLINK file set comprised of .bed, .bim and .fam files
DB=ddPCR_data_set_unrelated

# Filter variants not in chr 1, with any level of missingness or MAF<0.2
plink --bfile ${DB} --allow-no-sex --chr 1 --geno 0 --maf 0.2 --make-bed \
    --out ${DB}_filtered

# Make IBS-based distance matrix in square format
plink --bfile ${DB}_filtered --distance square 1-ibs --out ${DB}_filtered
```

The resulting matrices were used to generate phylogenetic trees following the distance-based tree-building algorithm Neighbor-Joining with function `nj` of the `ape` package (Paradis & Schliep, 2019). Trees were then manually rooted with the function `root` of the `tidytree` package (Yu, 2022) by the node representing the oldest split in the human phylogeny, as determined by **Figure 1a** of Duda and Zrzavý (2016). In the **ddPCR data set**, the rooting node was that separating Khwe and San (Ju/'hoansi or Khomani, as per HDGP naming) from all other groups. In the **read-depth data set**, which doesn't include Khwe people, the selected node was that basal to all San. Both trees were ultrametricized with function `chronos` of the `ape` package using default options.

```r
# Read 1-IBS matrix and format for later use
IBS_mdist <- read_table("ddPCR_data_set_unrelated_filtered.mdist", col_names=FALSE)
IBS_mdist.id <- read_table("ddPCR_data_set_unrelated_filtered.mdist.id", col_names=FALSE)
rownames(IBS_mdist) <- IBS_mdist.id$X2
colnames(IBS_mdist) <- IBS_mdist.id$X2

# Reorder IBS matrix to match order in the dataframe containing ddPCR data
ddPCR_data_set <- ddPCR_data_set[match(rownames(IBS_mdist), ddPCR_data_set$sample),]

# Make tree from 1-IBS matrix with NJ algorithm
tree <- nj(as.dist(IBS_mdist))

# Give tree tip labels and node numbers to identify best node for rooting
tree$tip.label <- ddPCR_modeling_data$pop
tree$node.label <- as.character(1:tree$Nnode)

# Plot tree to visualize node for rooting
```

```
plot(tree, "fan", show.node.label=TRUE, use.edge.length=FALSE, align.tip.label=TRUE)

# Select node for rooting
node.tree <- as_tibble(tree)[as_tibble(tree)$label=="130", "node"][[1]] # Ex. node: 130

# Root tree
tree_rooted <- root(tree, node=node.tree, resolve.root=TRUE)

# Chronometricize tree
tree_chrono <- chronos(tree_rooted, model="correlated")
```

Phylogenetic trees of both data sets are accessible in **Figure 4** and **Figure S4**.

#### 6.5 Phylogenetic variance-covariance (VCV) matrix

VCV matrices were obtained from the phylogenetic trees with function `vcv.phylo` of the `ape` package. The matrices were then incorporated into the modeling function `brm` of `brms` package as the covariance structure of the individual-level random effect using the `gr(individual, cov=VCV)` syntax. This approach allowed us to model autocorrelation among individuals based on their phylogenetic relationship.

```
# Obtain VCV matrix from rooted phylogenetic tree
VCV <- vcv.phylo(tree_chrono)
```

#### 6.6 Principal component analysis

The unsupervised dimensionality reduction algorithm implemented by `PLINK` function `pca` was run on chromosome one markers after linkage disequilibrium filtering with function `indep-pairwise 50 10 0.8`.

```
# PLINK file set comprised of .bed, .bim and .fam files
DB=ddPCR_data_set_unrelated

# Keep variants in chr 1
plink --bfile ${DB} --chr 1 --out ${DB}_chr1

# Identify variants in LD: 50 kb window size, 10 kb step size, 0.8 r2 threshold
plink --bfile ${DB}_chr1 --indep-pairwise 50 10 0.8 --out ${DB}_LD_results_50_10_0.8

# Remove variables in LD (keep variables in prune.in file)
plink --bfile ${DB}_chr1 --extract ${DB}_LD_results_50_10_0.8.prune.in --make-bed \
    --out ${DB}_chr1_LD_filtered_50_10_0.8

# Run PCR
plink --bfile ${DB}_chr1_LD_filtered_50_10_0.8 --pca \
    --out ${DB}_chr1_LD_filtered_50_10_0.8
```

We plotted the proportion of variance explained by each PC and determined the number of PCs to retain using the ‘elbow method’, which identifies the point at which additional components contribute diminishing returns in explained variance. The number of PCs included in the modeling of each data set was therefore variable.

### 7 Temporal signals in West Eurasia

#### 7.1 Supervised ADMIXTURE

Supervised `ADMIXTURE` (Alexander et al., 2009) was run as described below, where `FILE` is the name of a plink fileset containing all source individuals as well as the 533 ancient Europeans. The individuals selected to represent each of the three source populations were selected from the AADR data set and are listed in **Table 4**.

```
# Supervised ADMIXTURE with 3 sources
admixture --cv=10 -j3 --supervised ${FILE}.bed 3
```

**Table 4**. Individuals representative of the five test/source populations.

| Test population | Included individuals (names as in AADR) |
|---|---|
| Anatolian farmers | I1580, I1581, I1583, I0707, I0708, I0709, I0736, I0744, I0745, I0746, I1096, I1097, I1098, I1099, I1100, I1101, I1102, I1103, I1579, I1585, Bar31, Bar8, I11949, Bar25, I1583, I1583 |
| Western hunter-gatherers | Loschbour, Villabruna |
| Yamnaya pastoralists | I0443, I11446, I1917, I2105, I7489, I0357, I0370, I0429, I0439, I0441, I0444, Damgaard2018Yamnaya, MJ-06, RK1001, RK1007, SA601, ZO2002, RISE240, RISE546, RISE547, RISE550, RISE552, Bul4, I18794, I18801_d, I0438, I3141, I0231, BOY001 |

### 8 Out of Africa dispersal

#### 8.1 Geographical distances between non-Sub-Saharan populations and East Africa

Goegraphical distances between each non-Sub-Saharan African population and East Africa were calculated avoiding large water bodies to better represent likely migration routes. Several waypoints were defined (e.g., Arabian peninsula, Sunda peninsula, Bering straight) to facilitate the calculation of migration paths (see **Figure S5** for a graphical representation of all distances). The calculations were performed with function `distGeo` from the `geosphere` package (Hijmans, 2024), which allows the computation of distances for angular locations (e.g., longitude and latitudes on the globe).

```
# Migration waypoints
migration_origin <- c(39.5, 9.0)
arabian_peninsula <- c(38.5, 35.0)
turkey <- c(27.0, 38.0)
thailand <- c(95, 25)
singapore <- c(103, 1)
pakistan <- c(68, 28)
northern_china <- c(125, 50)
central_alaska <- c( -147.7, 64.8 )
bering_russia <- c( -171.0, 65.7 )
washington_center <- c(-120.5, 47.5)

# FUNCTION: calculate geographical distance between specified waypoints
```

```r
compute_path_dist <- function(waypoints) {
  total_distance <- 0
  for (i in 1:(length(waypoints)-1)) {
    # Add distances between all waypoints
    total_distance <- total_distance + distGeo(waypoints[[i]], waypoints[[i + 1]])}
  # Transform to kilometers
  return(total_distance/1000)}

n_min=5   # Min number of inds per pop

# Select data set to analyse
OOA_dataset <- RD_metadata %>%
  filter(region != "AFR") %>% # Keep non-Africans
  group_by(pop) %>%
  filter(n() >= n_min) %>% # Keep pops with  5 inds
  ungroup()

# Get population coordinates
OOA_pop_coords <- OOA_dataset %>%
  group_by(pop) %>%
  summarise(lon = first(longitude),
            lat = first(latitude),
            region = first(region))

# Calculate distance between East Africa and each population
OOA_geo_distances <- OOA_pop_coords %>%
  rowwise() %>% mutate(
    OOA_pop_coords = list(c(lon, lat)), # Pop coordinates
    migration_distance_km = case_when(  # Calculate OOA distances, varying routes:

      # AMERICA
      region=="AMR" ~
        compute_path_dist(list(migration_origin, arabian_peninsula, pakistan,
                               northern_china, bering_russia, central_alaska,
                               washington_center, OOA_pop_coords)),

      # OCEANIA
      region=="OCN" ~ compute_path_dist(list(migration_origin, arabian_peninsula,
                                             pakistan, thailand, singapore,
                                             OOA_pop_coords)),

      # SOUTH ASIA
      region=="SA" ~ compute_path_dist(list(migration_origin, arabian_peninsula,
                                            pakistan, OOA_pop_coords)),

      # EAST ASIA
      region=="EA" ~
        compute_path_dist(list(migration_origin, arabian_peninsula, pakistan,
                               thailand, OOA_pop_coords)),

      # CENTRAL ASIA and SIBERIA
      region=="CAS" ~
        compute_path_dist(list(migration_origin, arabian_peninsula, OOA_pop_coords)),
```

```
    # MIDDLE EAST
    pop %in% c("Druze","Palestinian","Mozabite","Bedouin") ~
      compute_path_dist(list(migration_origin, arabian_peninsula, OOA_pop_coords)),

    # WESTERN EURASIA
    region=="WEA" ~
      compute_path_dist(list(migration_origin, arabian_peninsula, turkey,
                             OOA_pop_coords))

  )) %>% ungroup()
```

#### 8.2 Measures of diversity at the *AMY1* locus

All populations with less than five individuals were removed from this analysis due to the high uncertainty that the sampled *AMY1* CN values were representative of the true population values. Nevertheless, the varying sample sizes for the remaining populations (5-25) could very well influence the statistics of diversity and therefore a rarefaction approach was followed to minimize the effects of disparate sample sizes. In this downsampling approach, we generated 500 replicates of five individuals each and averaged the diversity statistics obtained across replicates. Results presented in **Figure S5** include correlations between out of Africa distance and diversity statistics for both the downsampled and original populations.

```
# Calculate diversity metrics (no rarefication)
OOA_diversity_stats <- OOA_dataset %>%
  group_by(pop) %>%
  summarise(
    n = n(),
    avg_AMY1 = mean(AMY1_rd),
    SD_AMY1 = sd(AMY1_rd),
    Var_AMY1 = var(AMY1_rd),
    CV_AMY1 = sd(AMY1_rd)/mean(AMY1_rd),
    Range_AMY1 = diff(range(AMY1_rd)),
    UniqueValues_AMY1 = length(unique(AMY1_rd)),
    ShannonEntropy_AMY1 = calc_shannon(AMY1_rd),
    .groups = "drop")

# Calculate diversity metrics (with rarefication)
OOA_diversity_stats_RAREFIED <- OOA_dataset %>%
  group_by(pop) %>%
  do({
    results <- replicate(500, {   # Num of replicates
      samp <- sample_n(., n_min)  # Each replicate based on n_min individuals
      data.frame(
        n = n_min,
        avg_AMY1 = mean(samp$AMY1_rd),
        SD_AMY1 = sd(samp$AMY1_rd),
        Var_AMY1 = var(samp$AMY1_rd),
        CV_AMY1 = sd(samp$AMY1_rd)/mean(samp$AMY1_rd),
        Range_AMY1 = diff(range(samp$AMY1_rd)),
        UniqueValues_AMY1 = length(unique(samp$AMY1_rd)),
        ShannonEntropy_AMY1 = calc_shannon(samp$AMY1_rd))},
      simplify = FALSE)
    bind_rows(results) %>%
```

```
    summarise(across(everything(), mean))}) %>%
  ungroup()
```

### 9 Measures of phylogenetic signal

Several tests of phylogenetic signal were conducted independently of the main modeling framework to explore the properties of *AMY1* CN (i.e., the modeled variable) and its distribution along the human phylogeny. All tests were run with the `phylosignal` package (Keck et al., 2016), and consist of five statistical measures of phylogenetic signal, phylogenetic correlograms and local indicators of phylogenetic association (LIPA).

First, a `comparative data` R object was prepared to combine the phylogenetic tree and data frames already generated, and be used in the `phylosignal` tests:

```
# PREPARE OBJECT:
# AMY1 CN + POPULATION METADATA to be combined with PHYLOGENETIC TREE
tree <- comparative.data(phy=tree_chrono,
                         data=RD_modeling_data,
                         names.col="sample", vcv=TRUE, na.omit=FALSE)
# Create column with sample names
tree$data$sample <- phy$phy$tip.label
```

#### 9.1 Phylogenetic signal

Using the `phyloSignal` function with 1,000 permutations, we computed five complementary statistical measures of global phylogenetic signal: *Blomberg's* $K$ and $K*$ (Blomberg et al., 2003), *Pagel's* $\lambda$ (Pagel, 1999), *Abouheif's Cmean* (Abouheif, 1999), and *Moran's* $I$ (Gittleman & Kot, 1990). Each of these global metrics quantifies phylogenetic signal under different assumptions about trait evolution (see Münkemüller et al. (2012) for a detailed review).

*Blomberg's* $K$ and $K*$, as well as *Pagel's* $\lambda$ , assume a Brownian ('random walk') motion model of trait evolution and assess the extent to which trait variation is explained by phylogenetic relatedness under that model. On the other hand, *Moran's* $I$ and *Abouheif's Cmean* are autocorrelation indices, not based on an evolutionary model, and evaluate spatial autocorrelation in trait values across the tree. Together, these measures provide a robust assessment of whether *AMY1* CN exhibits any phylogenetic structure.

```
# Phylogenetic signal
phyloSignal(phylo4d(tree, data["AMY1_CN"]), reps=1000)
```

#### 9.2 Phylogenetic correlogram

To complement the global measures of phylogenetic signal and reveal its scale, we constructed a phylogenetic correlogram using the `phyloCorrelogram` function with default options. This method assesses the correlation of *AMY1* CN values between individual pairs across a range of phylogenetic distances and provides a graphical representation of how trait similarity is structured along the phylogeny. Positive correlations indicate that individuals within a given phylogenetic distance tend to have similar trait values, whereas negative correlations suggest dissimilarity in trait values among individuals at that phylogenetic distance. A commonly observed pattern in traits that evolve randomly across a phylogeny is positive correlation at shorter phylogenetic distances, gradually declining to zero or negative correlation at larger phylogenetic distances.

```
# Phylogenetic correlograms
phyloCorrelogram(phylo4d(tree, data["AMY1_CN"]), trait="AMY1_CN")
```

##### 9.3 Local indicators of phylogenetic association

The `lipaMoran` function was used to perform one-sided LIPA tests with 1,000 permutations. This method, based on the *Moran's I* statistic, evaluates each individual tip in the phylogeny to determine whether its trait value is significantly more similar or dissimilar to those of its phylogenetic neighbors than expected under a null model of trait evolution. This allows for the identification of localized clusters of similar *AMY1* CN values as well as potential outliers with divergent trait values.

To explore complementary patterns of local phylogenetic structure, the analysis was conducted twice using contrasting alternative hypotheses. The first test used the 'greater' alternative hypothesis, which detects clusters of similar values (either high or low *AMY1* CN) and is particularly suited to identifying regions of the tree where trait values are conserved. The second test used the 'less' alternative, which evaluates whether an individual's *AMY1* CN trait value is significantly dissimilar from its neighbors and can therefore reveal local trait divergence, highlighting evolutionary outliers or transitions where trait values shift markedly across closely related individuals.

```
# Local indicators of phylogenetic association
lipaMoran(phylo4d(tree, data["AMY1_CN"]), reps=1000, alternative="greater")
lipaMoran(phylo4d(tree, data["AMY1_CN"]), reps=1000, alternative="less")
```
